# Chromosome segregation synchrony in *S. pombe* is noise-limited and arises without positive feedback

**DOI:** 10.64898/2026.03.09.710577

**Authors:** Wendi Williams, Kien Phan, Jing Chen, Stefan Legewie, Julia Kamenz, Silke Hauf

## Abstract

Anaphase is a key cell cycle transition that ensures faithful genome inheritance. At anaphase onset, sister chromatids separate abruptly and synchronously upon activation of the protease separase. Major cell cycle transitions often involve positive feedback, which contributes to their abruptness and irreversibility; however, whether such feedback is required for anaphase remains unclear. Here, we analyze sister chromatid separation dynamics in fission yeast using high-resolution live-cell imaging and computational modeling. We find that anaphase synchrony relies on fast degradation of the separase inhibitor securin, but does not require separase-mediated positive feedback. Hence, sister chromatid separation, being inherently irreversible, may be one of the few major cell cycle transitions that can proceed without positive feedback. A stochastic model fitted to the data revealed that separation synchrony is limited by stochasticity resulting from small-number effects. Together, these results support a feedback-independent mechanism for anaphase onset and identify molecular noise as a fundamental constraint on its temporal precision.

## Introduction

The sudden, synchronous splitting of sister chromatids at anaphase is visually one of the most striking transitions in the cell cycle. Until anaphase, sister chromatids are held together by cohesin, a large protein complex that can topologically encircle two sister chromatids (i.e., create “cohesion”) and also influences global chromosome architecture by creating DNA loops (Uhlmann, 2025; Ochs and Gerlich, 2026; Nasmyth and Haering, 2009; Makrantoni and Marston, 2018). Cohesin is enriched at centromeric regions but also found along chromosome arms (Tomonaga et al., 2000; Schmidt et al., 2009; Watanabe, 2005). The establishment of cohesion during DNA replication ensures the proper attachment of sister chromatids to opposite poles of the mitotic spindle, a prerequisite for the equal distribution of genetic material to the two daughter cells (Tanaka et al., 2000; Sonoda et al., 2001). Sister chromatid cohesion is irreversibly lost during anaphase due to the proteolytic cleavage of the cohesin subunit Scc1 (Rad21 in fission yeast) by the protease separase (Uhlmann et al., 1999, 2000; Tomonaga et al., 2000; Hauf et al., 2001; Yu et al., 2025). Scc1/Rad21 cleavage releases the topological entrapment of sister chromatids, allowing them to move to opposite ends of the cell (Uhlmann et al., 2000; Pauli et al., 2008; Oliveira et al., 2010). Separase activation depends on the ubiquitination and subsequent degradation of its inhibitor, securin, mediated by the anaphase-promoting complex (APC/C) and the proteasome (Funabiki et al., 1996; Ciosk et al., 1998; Peters, 2006).

Across various organisms, chromosome separation occurs within a narrow time window, even though securin degradation proceeds much more slowly. For example, in human cells, complete securin degradation takes about 20 minutes, whereas the segregation of more than 40 chromosomes is completed within just 1 to 2 minutes (Hagting et al., 2002; Armond et al., 2019; Sen et al., 2021). Similar patterns are observed in mouse oocytes (McGuinness et al., 2009; Thomas et al., 2021) and budding yeast (Lyons and Morgan, 2011; Lu et al., 2014). To explain the abrupt onset of sister chromatid separation, it has been proposed that separase activity increases in a switch-like manner (Holt et al., 2008; Shindo et al., 2012; Yaakov et al., 2012; Hellmuth et al., 2014). Supporting this idea, cohesin cleavage has been observed to rise sharply just before sister chromatids separate (Shindo et al., 2012; Yaakov et al., 2012). Such a switch-like increase in separase activity could result from positive feedback regulation, a common feature of major cell cycle transitions (Kapuy et al., 2009; Ferrell, 2013). In budding yeast, a positive feedback loop was proposed in which separase enhances securin degradation by indirectly promoting dephosphorylation of securin at Cdk1-dependent sites and accelerating securin ubiquitination (Holt et al., 2008). However, subsequent studies found that non-phosphorylatable and wild-type securin are degraded at similar rates, and that the rate of cohesin cleavage is not strongly affected by expression of non-phosphorylatable securin (Yaakov et al., 2012; Lu et al., 2014). In human cells, PP2A-mediated dephosphorylation of separase-bound securin appears to decelerate rather than accelerate its degradation (Hellmuth et al., 2014; McGuinness et al., 2009). Thus, it remains unclear whether separase-mediated feedback is physiologically important and functionally conserved across eukaryotes.

To investigate the mechanisms behind sister chromatid separation synchrony, we implemented live-cell imaging with high temporal resolution in fission yeast. Fission yeast makes for a good model, as it has regional centromeres enriched in cohesin, similar to metazoan cells, only has three chromosomes that differ in size, and anaphase can be perturbed genetically or pharmacologically (Tong et al., 2019; Schmidt et al., 2009; Hayles and Nurse, 2017). Using live-cell microscopy, we quantified the timing and kinetics of sister chromatid separation and securin degradation in wild-type cells and analyzed how different perturbations alter these dynamics. We find that the synchrony of sister chromatid separation correlates with the speed of securin degradation. Combining these quantitative results with computational models suggests that positive feedback is not a necessary prerequisite for synchronous sister chromatid separation, and that instead the separation dynamics are explainable purely from stochastic separase-mediated cohesin cleavage. Fitting a stochastic model of cohesin cleavage to our experimental results reveals that synchrony is naturally limited by molecular noise.

## Results

### Sister chromatids separate within a narrow time window—but not with perfect synchrony

In fission yeast, securin (Cut2) degradation is completed in about 4 minutes, and chromosome I splits about 2 minutes after the onset of securin degradation (**Fig. S1A – D**) (Kamenz and Hauf, 2014). To analyze the time window in which the three chromosomes separate, we used live-cell imaging with high temporal resolution (3.5–7 sec) and monitored fluorescent fusion proteins recruited to the centromeres of the three fission yeast chromosomes (**Fig. 1A,B**). On chromosome II, LacI-GFP was targeted to a position close to the centromere with *lacO* repeats (cen2-GFP); on chromosome I and III, TetR-tdTomato was targeted to a position close to the centromere with *tetO* repeats (cen1- and cen3-tdTomato) (Sakuno et al., 2009; Straight et al., 1996; Yamamoto and Hiraoka, 2003). We compared separation timing between chromosome I and II, or II and III and defined one chromosome as separating when sister chromatids started a sustained movement towards opposite spindle poles and ceased to move coordinately or convergently (**Fig. 1C,D**). We typically scored separation based on visual inspection of the time-lapse recording, but tracking of the separation trajectories confirmed that the visually assigned separation times align with the onset of rapidly increasing inter-centromere distance (**Fig. S1E)**.

**Figure 1.**
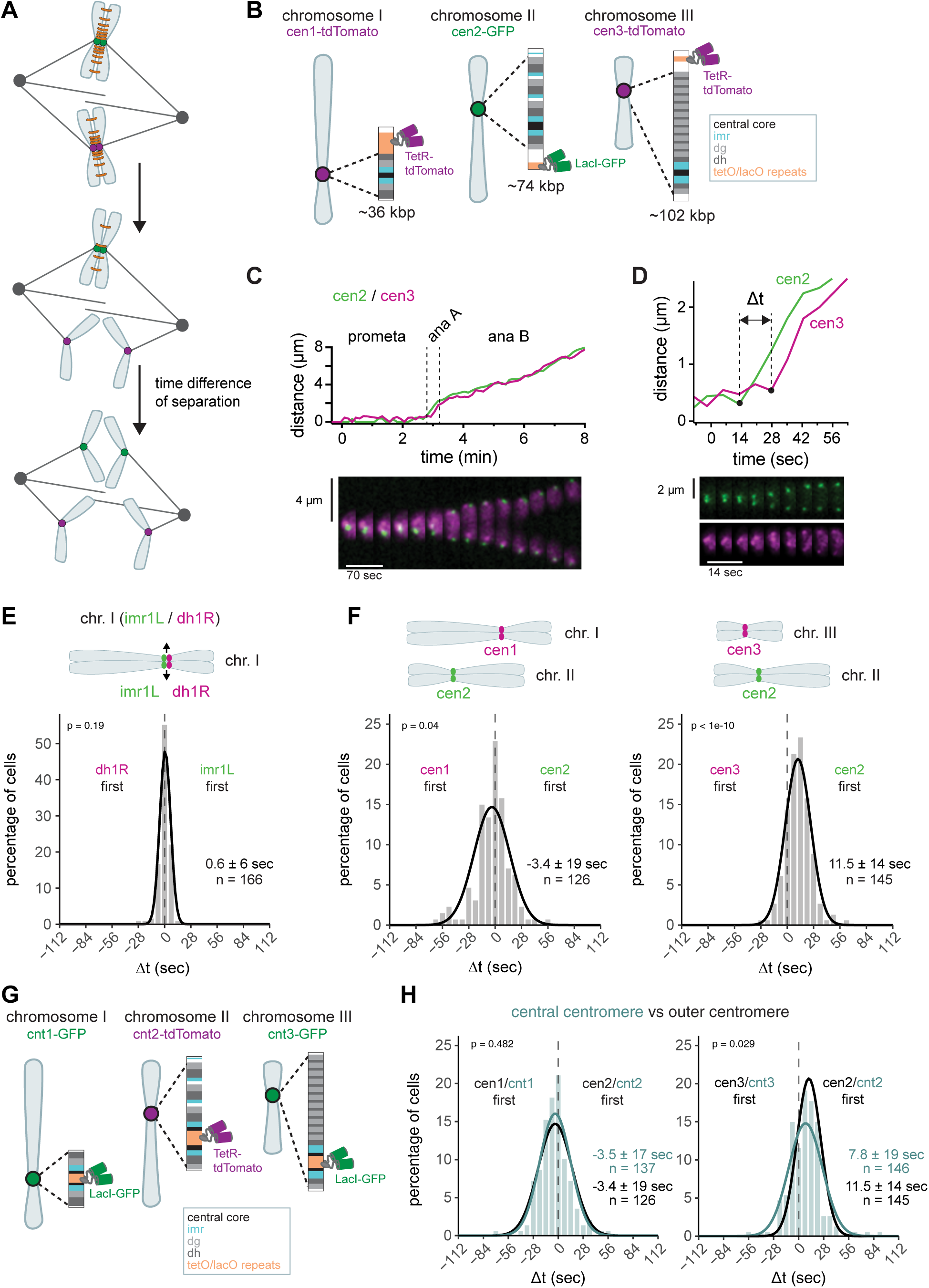
Centromeres segregate within a narrow time window but not with perfect synchrony. **(A)** Schematic depicting cohesin cleavage and subsequent sister chromatid separation; cohesin complexes in orange. **(B)** Fluorescent labeling of chromosomes close to their centromeric regions. The schematic depicts the localization of the tandem *tetO* or *lacO* repeats relative to the centromere, which comprises central core (cnt), innermost repeats (imr), and different numbers of dg/dh repeat pairs. Chromosome II was marked with *lacO*/LacI-GFP (Yamamoto and Hiraoka, 2003); chromosome I and III were marked with *tetO*/TetR-tdTomato (Sakuno et al., 2009 and this study). The lengths of the centromere regions (distance between furthest dh/dg repeats) are shown in kbp. **(C,D)** Sister centromere distance and corresponding kymograph for a strain with cen2-GFP and cen3-tdTomato marker. Same cell in (C) and (D). (D) shows how the time difference between two markers (Δt) is scored. **(E,F)** Frequency distributions and Gaussian fit (continuous lines) of the time difference (Δt) between the separation of two markers, either on the same chromosome (E), or on chromosomes I and II or chromosomes II and III (F). Mean ± standard deviation of the fit and number of cells are shown; p-value from one-sample t-test against zero. **(G)** Fluorescent labeling of chromosomes at the inner centromeric regions. Chromosome II was marked with *tetO*/tetR-tdTomato (Sakuno et al., 2009), whereas chromosome I and III were marked with lacO/LacI-GFP (Sakuno et al., 2009 and this study). **(H)** Cyan: Frequency distributions and Gaussian fit (continuous lines) of the time difference (Δt) between the separation of central centromere markers on chromosome I and II or chromosome II and III. The fitted Gaussian distributions of Δt using outer centromere markers (F) are shown in black for comparison. Mean ± standard deviation of the fit; n = number of cells; p-values from a two-sample Kolmogorov-Smirnov test.

The time difference of separation between two chromosomes (Δt, **Fig. 1D**) reflects the synchrony of anaphase; a 0 sec time difference indicates perfect synchrony. Indeed, monitoring two centromere markers on the same chromosome showed highly synchronous separation with a mean Δt of 0.6 sec (**Fig. 1E, S1F**). However, slight differences in separation timing between the markers were possible (standard deviation 6 sec), suggesting that separation of different parts of the centromere has a stochastic component. With markers on two different chromosomes, the mean of Δt was still close to, but distinct from zero, indicating a bias in the centromere of one chromosome separating before the other. Centromere 1 (cen1) split on average 3.4 sec before cen2, and cen2 split on average 11.5 sec before cen3 (**Fig. 1F**). However, this order was not absolute; despite cen2 separating on average 11.5 sec before cen3, cen3 separated before cen2 in about 23 % of the cells. The variations in Δt between individual cells, measured as the standard deviation of Δt, were 19 sec for cen1 versus cen2, and 14 sec for cen2 versus cen3, broader than for both markers on the same chromosome (6 sec) (**Fig 1E,F**, **Table S1**), indicating additional stochastic differences between the two chromosomes. The type of growth medium (rich versus minimal) did not have a significant influence on separation timing (**Fig. S1G**).

In budding yeast, it was initially proposed that chromosome segregation occurs in a strictly sequential order (Holt et al., 2008), but this was later attributed to one of the fluorescent tags (*tetO*/TetR-GFP) creating abnormally strong cohesion and delaying the separation of one chromosome (Lyons and Morgan, 2011). We therefore swapped the *tetO* and *lacO* markers for cen1 and cen2. While the presence of the *tetO* array may indeed very slightly bias separation towards later times, we did not observe any reversal in separation order, and the overall distribution remained highly similar (**Fig. S1H)**, indicating that separation timing is not greatly influenced by the nature of the fluorescent tag in our system.

The three centromere tags have different distances from the central core of the centromere where the kinetochore assembles and microtubules attach, and the marker on chromosome III was the furthest away from the central core (**Fig. 1B**). It was therefore possible that different separation timings reflect a “peeling apart” of the centromere region until the marker is reached. However, placing tags at the central core of the centromere, rather than the periphery, still resulted in similar Δt distributions (**Fig. 1G,H**). The bias for the centromere of chromosome II separating before that of chromosome III was slightly reduced but remained present (7.8 vs. 11.5 sec). This suggests that the centromeric region may separate as a single coherent unit. The chromosome arms, in contrast, separate distinctly later. A marker at the end of the long arm of chromosome II (Ding et al., 2004) split between 30 sec and more than 3 min after the chromosome II centromere (**Fig. S1I-K**). This is consistent with observations of the arms progressively peeling apart in both fission yeast and other eukaryotes (Ding et al., 2004; Paliulis and Nicklas, 2004; Renshaw et al., 2010; Chu et al., 2022). For the remainder of the experiments, we focused on centromere segregation as the earliest event of chromosome splitting.

Overall, these data quantify the synchrony of anaphase (∼15–20 sec standard deviation between two chromosomes) and indicate that centromere separation has a considerable stochastic component, so that the order of chromosome separation varies between cells.

### Chromosome segregation synchrony depends more on separase activity than on microtubule dynamics

The separation of sister chromatids is triggered by separase-mediated cohesin cleavage (Uhlmann et al., 2000); as such, synchrony likely requires rapid cohesin cleavage, which, in turn, is controlled by separase activation. Reducing separase activity, therefore, should make chromosome separation more asynchronous (reflected in a wider spread of Δt). To test this, we used the temperature-sensitive separase mutant *cut1-206* (Hirano et al., 1986) and followed sister chromatid separation at a semi-permissive temperature. When impairing separase activity, centromere separation for both chromosome pairs became significantly more asynchronous, more than doubling the standard deviation (**Fig. 2A, S2A-D**, **Table S1**). This is consistent with observations in budding yeast, where a separase mutant slowed down cohesin cleavage (Yaakov et al., 2012) and increased the variation in separation time between two chromosomes (Holt et al., 2008; Lyons and Morgan, 2011). We further observed a slower movement of centromeres towards the spindle poles in the separase mutant (**Fig. 2B, S2E**). This indicates a sustained requirement for separase activity beyond centromere splitting. Consistently, in budding yeast, the time between centromere separation and arm separation became longer when separase activity was reduced (Renshaw et al., 2010), which has been attributed to a requirement to remove remaining chromosome arm cohesin during early anaphase.

**Figure 2.**
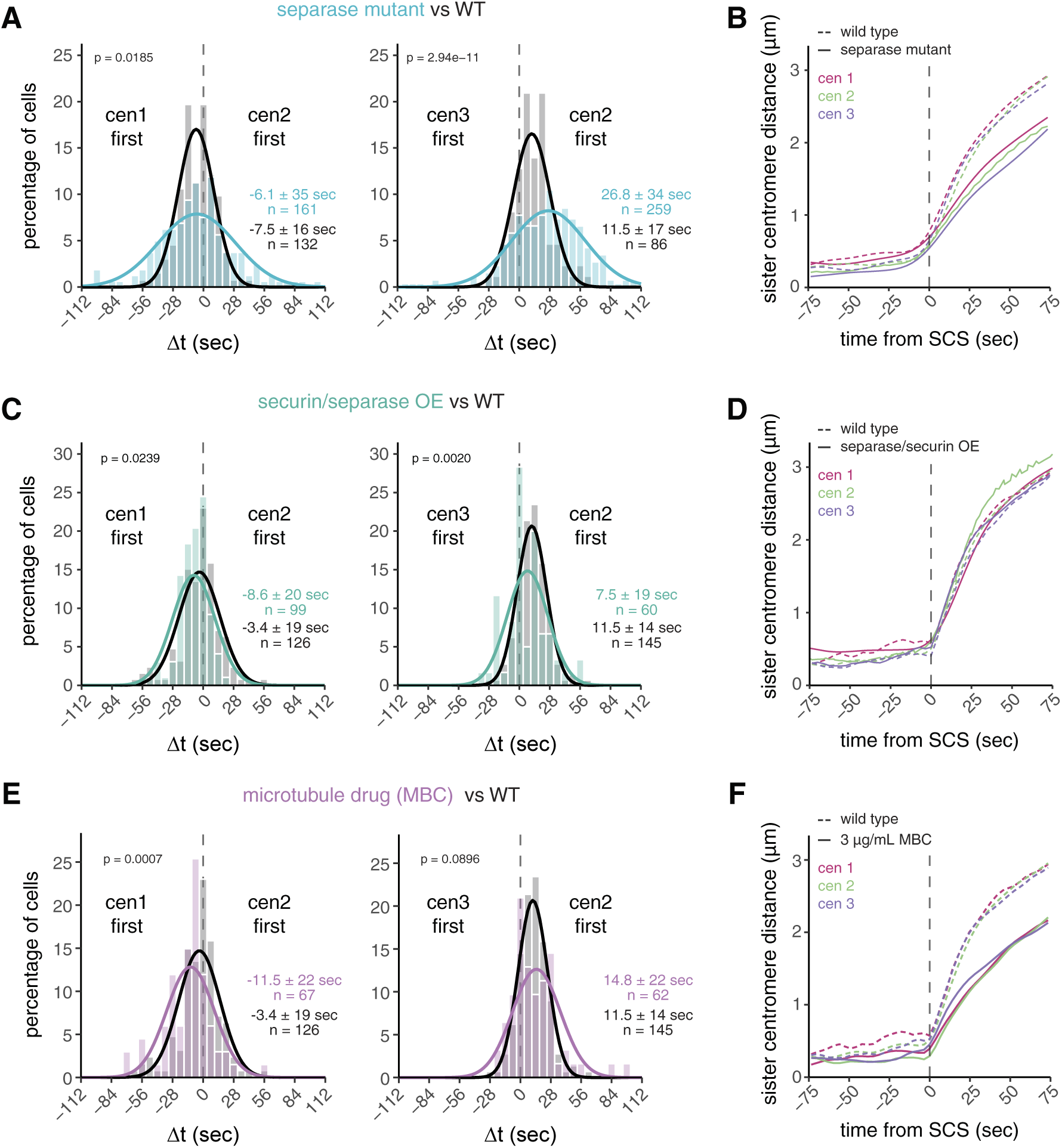
Separase activity has a larger influence on chromosome segregation synchrony than microtubule dynamics. **(A,C,E)** Frequency distributions and Gaussian fit (continuous lines) of the time difference between the separation of centromere 1 and 2 or centromere 2 and 3 for cells carrying the temperature-sensitive separase allele *cut1-206* (A), cells with overexpressed securin and separase (C), and cells exposed to 3 µg/mL of the microtubule drug MBC (E). The fitted Gaussian distributions of wild-type cells (black), imaged in the same medium as the experimental strain are shown for comparison (*cut1-206*: rich medium; securin/separase overexpression and MBC treatment: minimal medium). Mean ± standard deviation of the fit; n = number of cells; p-values from a two-sample Kolmogorov-Smirnov test. See Figs. S2-S4 for additional data. **(B,D,F)** Distances between sister chromatid pairs in wild-type cells (dashed lines) and cells carrying the temperature-sensitive separase allele *cut1-206* (B, solid lines), cells with separase and securin co-overexpression (D, solid lines), or cells exposed to 3 µg/mL MBC (F, solid lines). Colors correspond to the chromosome identity. Distances are aligned to the sister chromatid separation (SCS) time of the respective chromosome. At least 17 cells were averaged for each chromosome and genotype - exact cell counts are shown in Figs. S2-S4.

To address whether separase activity is limiting for chromosome separation synchrony in wild-type cells, we overexpressed separase. We co-overexpressed the inhibitor securin (**Fig. S3A,B**), both to prevent the potential lethality associated with uninhibited separase (Kamenz and Hauf, 2014) and to accelerate separase release since securin overexpression accelerates securin degradation kinetics (Kamenz et al., 2015). Neither securin overexpression alone (**Fig. S3C**) nor securin and separase co-overexpression (**Fig. 2C, S3D**) significantly reduced the variation of Δt. This suggests that, even though separase activity is important for a synchronous anaphase (**Fig. 2A**), additional factors limit anaphase synchrony. Sister chromatids still moved with the same speed in anaphase A in cells co-overexpressing securin and separase (**Fig. 2D, S3E,F**), suggesting that cleavage of remaining cohesin complexes on arms is not limited by separase availability.

Since these experiments suggested that mechanisms other than separase activity limit anaphase synchrony, we sought to determine whether forces exerted by microtubules of the mitotic spindle play a role. Microtubules undergo growth and shrinkage, and kinetochores attached to microtubules oscillate between the spindle poles, partially also influenced by motor proteins (VandenBeldt et al., 2006; Cheeseman and Desai, 2008; Walczak et al., 2010). Hence, the separation of centromeres may be delayed after cohesin cleavage until microtubules pull towards opposing poles. Indeed, in *Drosophila*, microtubule flux has been shown to contribute to anaphase synchrony (Matos et al., 2009). We used two perturbations to impair microtubule dynamics, the microtubule drug MBC and deletion of the kinesin-8, *klp5*; MBC is expected to weaken microtubules, *klp5Δ* is expected to stabilize microtubules and alter chromosome oscillations (Balestra and Jimenez, 2008; Garcia et al., 2002; West et al., 2002; Unsworth et al., 2008; Klemm et al., 2018). We titrated MBC to a concentration that still allowed proper chromosome segregation in almost all cells but slowed down anaphase centromere movement (**Fig. 2F, S4A,B**). Anaphase synchrony was slightly worsened by MBC – the standard deviation of Δt for cen1 vs. cen2 increased from 19 to 22 sec, and for cen2 vs. cen3 from 14 to 22 sec (p = 0.3970 and 1.92e-4, respectively, by Levene’s test) (**Fig. 2E**). However, this change was considerably less than that observed in the separase mutant, which—while having similarly slow centromere movement during anaphase—had standard deviations of 35 sec for cen1 vs. cen2 and 34 sec for cen2 vs. cen3 (p = 4.78e-9 and 1.17e-7, respectively, by Levene’s test) (**Fig. 2B,F**). Furthermore, anaphase synchrony was unchanged in *klp5Δ* cells (**Fig. S4C**), even though chromosomes were often misaligned at the onset of anaphase, as expected (**Fig. S4D**). Together, our data therefore suggest that microtubules play a minor role in *S. pombe* centromere separation synchrony compared to separase.

### No evidence for separase-mediated feedback on securin degradation to enhance chromosome segregation synchrony

As separase activity plays an important role in the synchrony of centromere separation, we considered possible mechanisms that allow for a rapid, sudden increase in separase activity. In budding yeast, a positive feedback loop has been proposed to contribute to synchrony, where separase enhances the degradation of its inhibitor, securin, by promoting the dephosphorylation of CDK1-dependent phosphorylation sites on securin (Holt et al., 2008). Persistent CDK1 activity (through expression of a non-degradable version of the cyclin Clb5) decreased sister chromatid separation synchrony. However, it is unclear whether this is caused by disruption of the proposed feedback loop or by the sustained CDK1 activity, independent of securin degradation.

We conducted a similar experiment in fission yeast and conditionally expressed a non-degradable version of the B-type cyclin Cdc13 (*ΔN-cyclin B*) at close to endogenous levels (Kamenz and Hauf, 2014). As a consequence, a fraction of cells was unable to divide the nucleus and exit mitosis, as is expected from sustained CDK1 activity (Yamano et al., 1996). Other cells, presumably with lower levels of non-degradable cyclin B, still underwent nuclear division and exited mitosis (**Fig. 3A,B**). The synchrony of sister chromatid separation in both classes of cells was comparable to that of wild-type cells (**Fig. 3A**, **Table S1**). This made it unlikely that fission yeast has a similar type of feedback loop as proposed for budding yeast. Consistent with the similar separation synchrony, the securin degradation rate also remained highly similar to that in wild-type cells when non-degradable Cdc13 was expressed (**Fig. S5A**) (Kamenz and Hauf, 2014), making it unlikely that separase affects securin degradation through CDK1. To test more broadly whether separase enhances securin degradation—independent of CDK1—, we expressed two versions of securin: wild-type securin tagged with GFP to monitor securin degradation, and inducible, untagged, non-degradable securin (ΔN-securin) to block separase activity. If separase enhances securin degradation, securin degradation should be slowed down when separase release is blocked by non-degradable securin. We confirmed this prediction by implementing a computational model for the key reactions with or without feedback (**Fig. 3C,D**; **Supplemental Material**). Experimentally, we found that the presence of non-degradable securin blocked nuclear division (**Fig. 3F**), indicating a failure of separase activity; yet, degradation of the wild-type version of securin was unaffected (**Fig. 3E,F, S5B**). Hence, separase activity does not accelerate securin degradation in fission yeast. We conclude that a positive feedback loop, where separase enhances its own activation by accelerating securin degradation, is not an integral part of the mechanism that promotes synchronous sister centromere separation in fission yeast.

**Figure 3.**
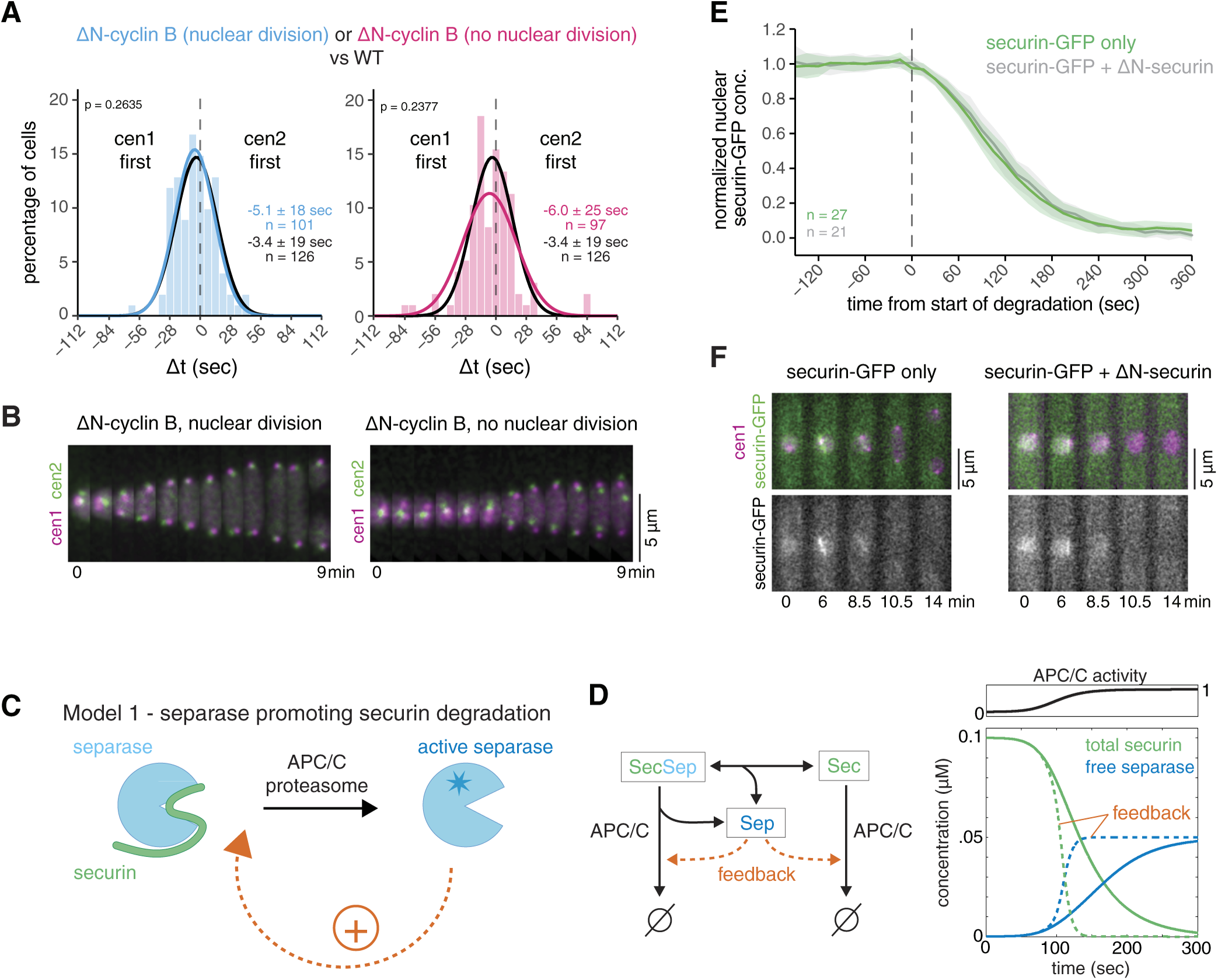
Separase-mediated feedback on securin degradation is unlikely. **(A)** Frequency distribution and Gaussian fit (continuous lines) of the time difference between the separation of centromere 1 and 2 after expression of non-degradable cyclin B (Cdc13), ΔN-cyclin B, in minimal medium. The Gaussian fit of wild-type cells grown under similar conditions (black) is shown for comparison. Mean ± standard deviation of the fit; n = number of cells; p-values from a two-sample Kolmogorov-Smirnov test. **(B)** Representative kymographs of sister chromatid separation in cells with cen1-tdTomato and cen2-GFP after expression of non-degradable cyclin B (Cdc13), ΔN-cyclin B, in minimal medium. The left panel depicts a cell which undergoes nuclear division and progresses through mitosis; the right panel displays a cell that fails to undergo nuclear division. **(C)** Schematic depicting a positive feedback loop, where separase accelerates its own activation by accelerating securin degradation. **(D)** Computational model for APC/C-mediated securin degradation and separase release with or without feedback of separase on securin degradation. Left side: diagram of the reactions present in the model. Right side: simulation with or without feedback (dashed and solid lines, respectively) assuming a sigmoidal increase of APC/C activity with time. See Supplemental Material for model details. **(E)** Securin-GFP degradation in wild-type cells (green) or in cells failing to separate their chromosomes after induction of non-degradable securin (Cut2), ΔN-securin (grey). Individual time courses are normalized and aligned to the start of securin degradation at t=0. Mean (line) ± standard deviation (shaded area) of the cell population; n = number of cells. **(F)** Representative kymographs of securin degradation in a wild-type cell (left panel) and in a cell after induction of untagged non-degradable securin (Cut2), ΔN-securin. In the latter cell, the nucleus fails to divide.

### No evidence for separase autoactivation to enhance chromosome segregation synchrony

A positive feedback loop would not necessarily need to act upstream to promote separase release from securin (**Fig. 3C**); it could also operate downstream, such as through separase autoactivation (**Fig. 4A**). For example, metazoan separase is known to cleave itself autocatalytically, which could alter its activity (Waizenegger et al., 2002; Zou et al., 2002; Chestukhin et al., 2003; Shindo et al., 2022). However, there is no clear evidence that cleavage enhances activity (Papi et al., 2005; Holland et al., 2007; Shindo et al., 2022), and separase auto-cleavage has not been observed in yeast (Hornig et al., 2002). Nevertheless, to analyze the possibility of downstream positive feedback, regardless of the mechanism, we simulated securin degradation and separase release with or without separase autoactivation in a computational model (**Fig. 4A,B**; **Supplemental Material**). The two model versions differ in their response to decreased APC/C activity. With or without feedback, decreased APC/C activity slows down securin degradation; however, without feedback, separase release becomes more gradual (**Fig. 4B, left graph)**, whereas with feedback, separase release remains abrupt but occurs later (**Fig. 4B, right graph, S6A**). If separase autoactivates, one would therefore expect that reduced APC/C activity delays sister chromatid separation but does not affect synchrony.

**Figure 4.**
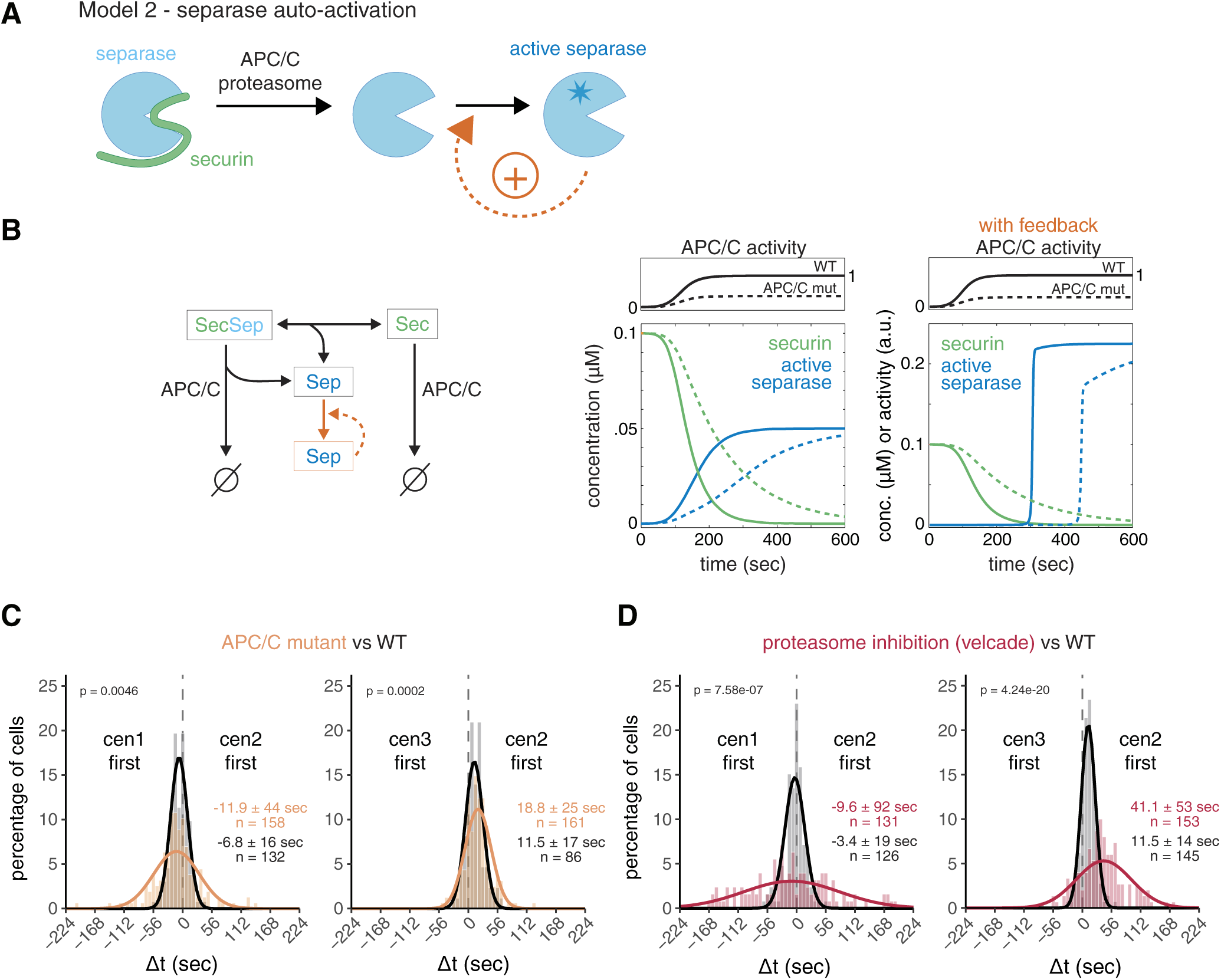
Separase auto-activation is unlikely. **(A)** Schematic depicting a positive feedback loop, where separase accelerates its own activation downstream of securin degradation. **(B)** Computational model for separase release mediated by APC/C-mediated securin degradation with or without separase auto-activation. Left side: diagram of the reactions present in the model. Right side: simulation for high or low APC/C activity, without or with feedback (separase auto-activation), assuming a sigmoidal increase of APC/C activity with time. See Supplemental Material for model details. **(C,D)** Frequency distributions and Gaussian fit (continuous lines) of the time difference between the separation of centromeres 1 and 2 or 2 and 3. Cells were carrying either a temperature-sensitive allele of the APC/C subunit Cut9 (cut9-665) and were grown in rich medium before imaging (C, orange), or were grown in minimal medium and treated with 100 µM of the proteasome inhibitor velcade (bortezomib) 30 min prior to imaging (D, red). The fitted Gaussian distribution of wild type cells grown under similar conditions and without inhibitor is shown for comparison (black). Mean ± standard deviation of the fit; n = number of cells; p-values from a two-sample Kolmogorov-Smirnov test.

To test this, we employed a temperature-sensitive allele of the APC/C subunit APC6 (*cut9-665*) grown at semi-permissive temperature (Hirano et al., 1986). Securin degrades slower in this mutant, as expected (**Fig. S6B,D**). Inconsistent with separase autoactivation, sister chromatid separation became less synchronous (standard deviations of Δt 44 and 25 sec) (**Fig. 4C, Table S1**). To corroborate this result, we slowed down securin degradation with a second method—by partial inhibition of the proteasome with velcade (bortezomib) (Takeda et al., 2011). This yielded even slower securin degradation than in the APC/C mutant (**Fig. S6C,D**) and further decreased the synchrony of sister chromatid separation (standard deviations of Δt 92 and 53 sec) (**Fig. 4D, S6E**, **Table S1**). Because of this pronounced effect of securin degradation kinetics on the synchrony of sister chromatid separation, separase autoactivation is unlikely.

### The dynamics of chromosome segregation can be reproduced by a basic stochastic model of cohesin cleavage

Since our results argue against separase-mediated positive feedback, we sought to determine if minimal features of separase regulation are sufficient to account for the sister chromatid separation dynamics in wild-type and mutant cells. To test this, we implemented a stochastic model of separase-mediated cohesin cleavage (**Fig. 5A; Supplemental Material**). We assumed that separase activity increases gradually over the period t until it reaches its maximal rate (k_max_), and that separase stochastically cleaves cohesin complexes on the three different chromosomes. Sister chromatid separation occurs when the number of cohesin complexes on one chromosome (N) has been reduced to the threshold number n. We do not necessarily assume that the threshold number n is zero, because, if the forces on a cohesin complex are large enough, it will let go of the bound DNA even without cleavage (Daum et al., 2011; Richeldi et al., 2024). We set boundaries for all parameters based on our data and the available literature (see **Table S4**) and fitted the model to the experimental data from wild-type cells, as well as all perturbations (separase and APC/C mutants, velcade, and MBC). We assumed (i) that the separase mutant changes separase activity, which we represent as a reduction in k_max_, (ii) that the APC/C mutant and velcade—by slowing down securin degradation—change the speed of separase activation, which we represent as a longer t, and (iii) that MBC—by lowering microtubule forces–reduces how many cohesin molecules can remain at the time of separation, which we represent as lower threshold number, n.

**Figure 5.**
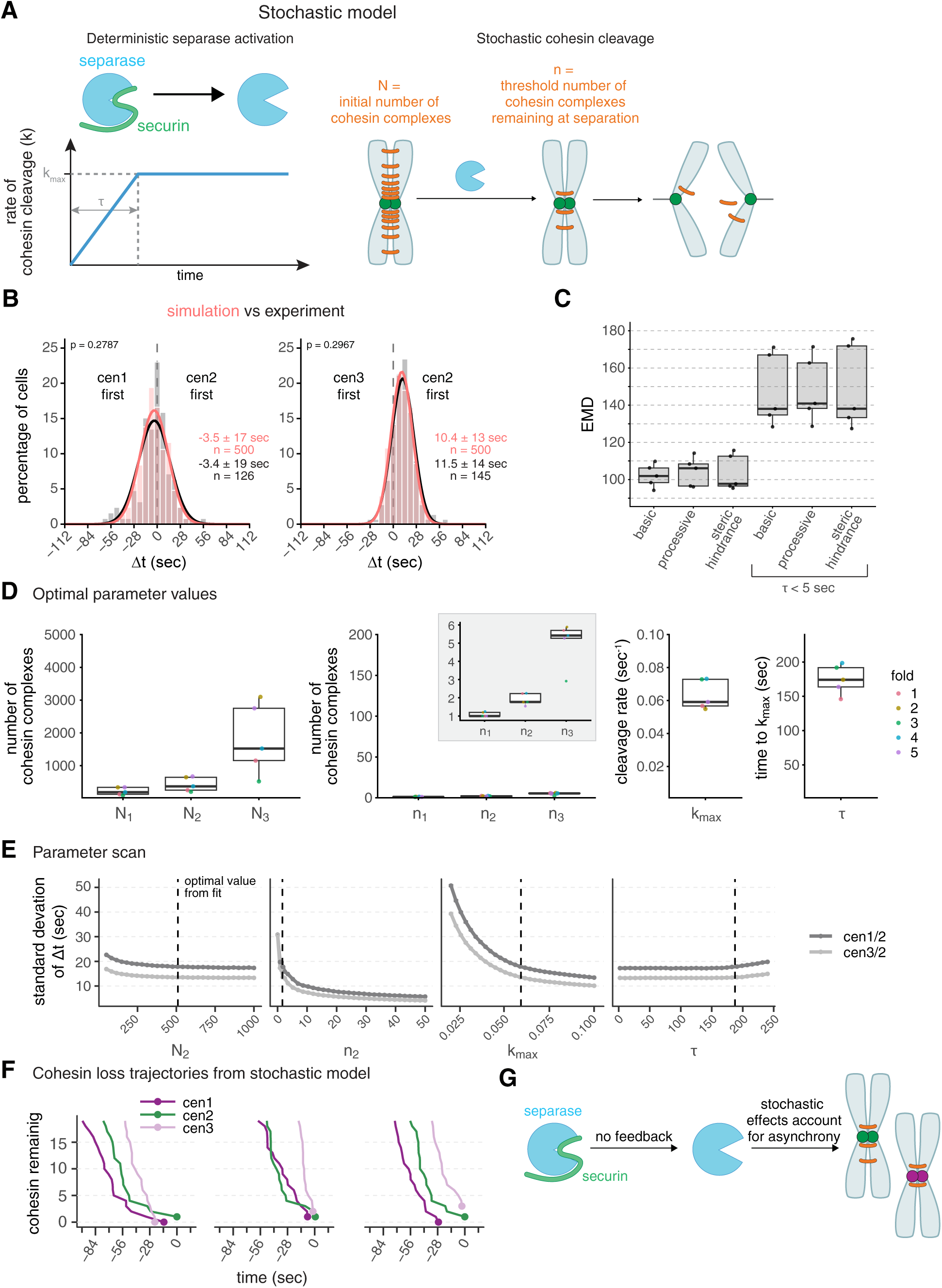
Small-number effects cause stochasticity in sister chromatid separation time. **(A)** Schematic of the stochastic model of separase-mediated cohesin removal. Separase activation (through securin degradation) is represented by a deterministic increase in the cohesin cleavage rate, k. The rate increases for a period τ, until it reaches its maximum value k_max_. Each chromosome initially has N cohesin complexes (N_1_, N_2_, N_3_, for chromosome I, II, and III, respectively), which are randomly cleaved by separase. Chromosomes separate once the number of remaining cohesins falls below a threshold n (n_1_, n_2_, n_3_). See Supplemental Material for model details. **(B)** Frequency distributions and Gaussian fit (continuous lines) of time differences between the separation of centromere 1 and 2 or centromere 2 and 3, either from a simulation using optimal parameter values of the basic model (salmon) or determined experimentally in wild-type cells (gray/black, same data as in Fig. 1F). Mean ± standard deviation of the fit; n = number of cells; p-values from Earth Mover’s Distance (EMD). **(C)** Goodness-of-fit of the model variants measured by Earth Mover’s Distance (EMD, see Supplemental Material), with and without constraining τ at low values (τ <5 sec). Points indicate the validation EMD values obtained in the five-fold cross-validation method (see Supplemental Material); boxplots show median, interquartile range and range. Basic, basic model with time-dependent cleavage rate alone; processive, model with bursts of cohesin cleavage; steric hindrance, model in which only a surface fraction of cohesin is available for cleavage and inner layers of cohesin are progressively exposed (see Supplemental Material). **(D)** Optimal parameter values obtained by fitting the basic model to the experimental data. Points indicate individual fold values (see Supplemental Material); boxplots show median, interquartile range, and range; the y-axis spans the parameter range tested, except for the inset showing n_1_, n_2_, n_3_. **(E)** One-at-a-time (OAT) sensitivity analysis of the basic stochastic model. Each parameter was varied individually across its allowed range while all other parameters were held fixed at their optimal values obtained by fitting. 10 batches of 10,000 simulations were performed for each parameter point. Each batch yielded a mean and standard deviation of the separation time differences (Δt) between centromere 1 and 2, and centromere 2 and 3. Plot shows mean of the Δt standard deviation +/- SEM across 10 batches. Dashed vertical lines indicate the optimal parameter values. **(F)** Stochastic cohesin loss trajectories near the time of separation in the basic model. Simulated cohesin counts for chromosomes 1, 2, and 3 are shown for three representative simulations, aligned to t=0 at the final separation event. Lines indicate cohesin loss over time; points mark the moment of separation for each chromosome. **(G)** Schematic summarizing the results: separase becomes active upon securin degradation without positive feedback. When a small number of cohesin complexes suffices to maintain cohesion, stochasticity of the final cleavage events dominates the timing of sister chromatid separation, leading to asynchrony and no clear order between chromosomes.

This model was able to reproduce the trends in sister chromatid separation synchrony across all experimental conditions (**Fig. 5B, S7A**), providing additional evidence that feedback regulation is not necessary. In fact, forcing t to be very short, which mimics the separase activity dynamics with separase auto-activation (**Fig. 4B**), makes the fit worse (**Fig. 5C**, basic vs. basic τ < 5 sec; **S7A**). To further probe whether adding regulatory mechanisms to the basic model would improve the fit, we considered two plausible variations: (i) processive separase action, where not one but multiple cohesin molecules are removed by each separase action, and (ii) steric hindrance, where initially not all cohesin is available to separase and only becomes available progressively (**Supplemental Material**). While both these additions provide good fits, they do not substantially improve the fit (**Fig. 5C**). Hence, a basic stochastic model of separase-mediated cohesin cleavage is sufficient to explain the experimentally observed dynamics of sister chromatid separation. In contrast, assuming separase autoactivation (τ < 5 sec) produces dynamics that are significantly less consistent with the experimental data (**Fig. 5C, S7A**).

### Small-number effects limit separation synchrony

To identify the main determinants of separation synchrony, we inspected the value of the parameters obtained by fitting (**Fig. 5D**) as well as the model’s sensitivity to changes in individual parameters (**Fig. 5E, S7B,C**). The optimal value for the starting number of cohesin molecules on each chromosome (N) was highest on chromosome 3, lower on chromosome 2, and yet lower on chromosome 1 (**Fig. 5D**). This is explained by the fact that the ratio of N between two chromosomes influences the mean Δt (**Fig. S7B**). Hence, the different starting numbers account for the preferential order observed (chromosome 1 being slightly more likely to separate before chromosome 2, and chromosome 2 more often separating before chromosome 3 than vice versa, **Fig. 1F, S1G**).

Two parameters, n and k_max_, strongly influence the synchrony of sister chromatid separation (**Fig. 5E**). Most notably, the value of n (threshold number of cohesin molecules remaining at the time of separation) that we obtained by fitting was at the lowest edge of the allowed parameter range (**Fig. 5D**). In this low-number regime, the timing of the final few cohesin cleavage events is stochastic, which decreases synchrony in the separation time and randomizes the order of separation (**Fig. 5E,F, S7C**). Synchrony is also decreased by low separase activity (**Fig. 5E**), because the timespan between stochastic degradation events and therefore between final separation events increases. Together, these results indicate that small-number effects place a natural limit on the synchrony of sister chromatid separation. In other words, if a small number of cohesin molecules is sufficient for cohesion (which is plausible, see Discussion), asynchrony in chromosome separation is expected.

## Discussion

Anaphase is visually one of the most striking steps of the cell cycle. When mitosis is imaged in live cells, the splitting of chromosomes into their sister chromatids occupies only a short period within mitosis and appears abrupt and synchronous. However, imaging at high temporal resolution, as we did here using fission yeast, and has been done in human cells (Armond et al., 2019; Sen et al., 2021; Harrison et al., 2021), reveals that sister chromatid separation is not perfectly synchronous and that there is significant variation in the segregation time of single chromosomes (**Fig. 1**). We propose that there is a natural limit to separation synchrony resulting from small-number effects.

### Separase-mediated positive feedback is not necessary to explain the dynamics of sister chromatid separation

In budding yeast, lower separase activity makes chromosome segregation more asynchronous (Holt et al., 2008; Lyons and Morgan, 2011) (**Fig. 2**), and a focus of prior studies has therefore been on separase regulation. Positive feedback loops are common in the regulation of cell cycle transitions and support both a rapid transition and irreversibility (Kapuy et al., 2009; Ferrell and Ha, 2014; Ferrell, 2013). It was therefore reasonable to assume that anaphase, as one of the major cell cycle transitions, makes use of positive feedback (Holt et al., 2008). Our results, however, argue against separase activity-enhancing positive feedback in anaphase regulation in fission yeast (**Figs. 3,4**). Anaphase is special among the cell cycle transitions as its irreversibility is inherent in its mechanics—the loss of chromosome cohesion cannot be reversed. Sister chromatid separation in anaphase may therefore be a major cell cycle transition that can dispense with positive feedback.

Other non-feedback mechanisms support rapid separase activation and cohesin cleavage, though (Kamenz and Hauf, 2017). Securin is a stoichiometric inhibitor of separase, present in excess over separase (Ciosk et al., 1998; Shindo et al., 2012; Kamenz et al., 2015; Hellmuth et al., 2014). Sequestration of separase by securin allows in principle for a rapid activation of separase once the excess securin pool is exhausted—known as inhibitor ultrasensitivity (Buchler and Cross, 2009; Ferrell and Ha, 2014; Legewie et al., 2008; Hopkins et al., 2017). In human and mouse cells, unbound securin is degraded before separase-bound securin (Hellmuth et al., 2014; Thomas et al., 2021), reinforcing that the free pool is exhausted first. Various post-translational modifications on securin, separase, and cohesin also support separase activity and separase-mediated cohesin cleavage (Alexandru et al., 2001; Yaakov et al., 2012; Li et al., 2017; Wang et al., 2024; Lianga et al., 2018; Yu et al., 2025). Even with such regulation in place, though, separase release will still slow down when securin degradation is slowed, consistent with our experimental results (**Fig. 4**). However, when we raised separase activity (**Fig. 2**), sister chromatid separation remained asynchronous. This suggests that once separase activation is sufficiently fast, increasing its activity no longer strongly improves synchrony. We propose a limit to separation synchrony that is determined by the nature of cohesin-mediated cohesion.

### Small-number effects as a natural limit to chromosome separation synchrony

Fitting a stochastic model of separase-mediated cohesin cleavage to our data revealed that the observed variation in chromosome separation order can be explained by the stochasticity that arises when a small number of cohesin molecules suffices for cohesion (**Fig. 5**). Is this plausible? Maybe yes. Studies that titrated cellular cohesin levels found that sister chromatid cohesion was maintained at levels as low as 13 % of wild-type cohesin in budding yeast (with lower levels not tested) or 22 % in Drosophila cells (Heidinger-Pauli et al., 2010; Carvalhal et al., 2018). More directly, the force required to rupture a fission yeast cohesin complex connecting two DNA molecules has been carefully measured *in vitro* (Richeldi et al., 2024) and was found to be in the same range as the force that a kinetochore-microtubule is thought to generate (Gudimchuk and Alexandrova, 2023; Nicklas, 1983; Akiyoshi et al., 2010; Volkov et al., 2013). Yeast kinetochores attach to 1–4 microtubules (Ding et al., 1993; Winey et al., 1995; O’Toole et al., 1999), suggesting that indeed a few cohesin molecules may suffice to maintain cohesion, consistent with the expectations from our stochastic model.

In metazoan cells, with a larger number of kinetochore microtubules, more cohesin complexes will be required to maintain cohesion (possibly in the order of ∼40 (Richeldi et al., 2024)). This may reduce the stochasticity in chromosome separation and establish a more predictable order determined by the difference in the initial amount of cohesin at the centromeres. Short-time-frame imaging in human cells found that separation of the first and last chromosome can be spaced by as much as 90 sec (Armond et al., 2019; Sen et al., 2021). However, individual chromosomes were not identified, and it therefore remains unclear how much this reflects a defined temporal order versus stochastic variation in separation times. Chromosome spreads of early anaphase chromosomes from metazoan cells suggest that chromosomes with smaller centromeres separate before those with larger centromeres (Vig, 1983). In addition, a distinct order in the separation of different mammalian chromosomes was also inferred from an observed inheritance of chromosome positions within the nucleus (Gerlich et al., 2003). Together, this suggests that the order of chromosome separation may be more pronounced in mammalian cells compared to fission yeast, possibly because small-number effects are minimized.

In summary, we propose that asynchrony in fission yeast sister chromatid separation, and potentially other cell types, arises because only a handful of cohesin molecules are left when cleaving one more leads to separation (**Fig. 5**). We base this on the good fit between our experimental data and a basic stochastic model of cohesin cleavage. In the spirit of Occam’s razor, using a simple model seems justified, but we acknowledge that stochastic variations in separation order could also result from factors not covered in our model. This includes stochastic variations in the amount of cohesin loaded onto single chromosomes, variation in centromere packing that alters access to cohesin, or variation in spindle forces. In *Drosophila* cells, for example, microtubule flux homogenizes the microtubule force across chromosomes and thereby contributes to chromosome segregation synchrony (Matos et al., 2009). To analyze the role of small-number effects, future experiments could make use of single-molecule imaging or test the effect of cohesin complexes that can resist higher force because DNA gates are covalently closed (Haering et al., 2008; Richeldi et al., 2024). It is often stated that synchronous chromosome separation is important for proper genome inheritance, but, overall, our results suggest that a small amount of separation asynchrony is unavoidable and is well tolerated.

## Materials and methods

### *S. pomb*e strains

All strains are listed in **Supplementary Table S2**. *S. pombe* strains with the following modifications and mutations have been described previously: *dh1L(cen1)-tdTomato*, *cnt1-GFP*, *cnt2-tdTomato* (Sakuno et al., 2009), *cen2-GFP* (Yamamoto and Hiraoka, 2003), *cut9-665* and *cut1-206* (Hirano et al., 1986), *cut2-GFP<<kanR*, securin overexpression (*natNT2<<Padh1(#6)-cut2-GFP<<kanR*), and the inducible version of non-degradable cyclin B (*leu1+<<Pnmt81-DN67-cdc13*) (Kamenz and Hauf, 2014). To fluorescently label the region close to the centromere of chromosome III, the plasmid pSR14 was used to integrate a ∼224x*tetO* array 556 bp 5’ of *meu27* following the previously described method (Rohner et al., 2008). In a similar manner, to fluorescently label the central centromere of chromosome III, the plasmid pSR13 was used to integrate a ∼248x*lacO* array 2,369 bp 3’ of the start of the central centromere (Rohner et al., 2008). For inducible expression of non-degradable securin (ΔN-Cut2), the coding sequence of amino acids 81 to 301 of Cut2 (Funabiki et al., 1996) was cloned into the pDUAL vector (Matsuyama et al., 2004) under the control of the *nmt81* promoter and integrated at the *leu1* locus. For deletion of endogenous *klp5*, an *hphMX6* resistance cassette was inserted using PCR-based deletion methods for *S. pombe* (Bähler et al., 1998). For separase overexpression, *cut1* was cloned into a pDUAL vector (Matsuyama et al., 2004) under the control of the *cut1* promoter and integrated at the *leu1* locus, or the endogenous *cut1* promoter was replaced with an *ark1* promoter.

### Cell cultures and live cell imaging

Prior to imaging, cells were cultured either in rich medium (YEA, yeast extract + adenine) or Edinburgh minimal medium (EMM) with the necessary supplements (Moreno et al., 1991). Cells were cultured at 30 °C except for the strains carrying the temperature-sensitive alleles *cut9-665* or *cut1-206*, which were cultured at 25 °C and incubated at 30 °C for 30 min prior to imaging at 30 °C. We used cells grown in EMM for the proteasome inhibition, because this led to a more effective inhibition, while the *cut9-665* and *cut1-206* phenotypes were more pronounced after growth in rich medium. The *nmt81* promoter was repressed by the addition of 16 µM thiamine to EMM and induced by transferring the cells into EMM without thiamine for 14–18 hours. Immediately prior to imaging, strains were transferred into EMM, except for the *cut9-665* strains and *cut1-206* strains, which were imaged in YEA. Cells were mounted in lectin-coated (35 µg/ml, Sigma L1395) culture dishes (8-well, Ibidi) and pre-incubated on the microscope stage at 30 °C for 30 min. For partial inhibition of the proteasome, Bortezomib (Velcade, LC Laboratories, B-1408) was added to a final concentration of 100 µM. For partial destabilization of microtubules, 3, 5, or 10 µg/mL of carbendazim (MBC) was added 20–30 minutes before imaging. Live cell imaging was carried out at 30 °C on a DeltaVision Core system (Applied Precision/GE Healthcare) equipped with a climate chamber (EMBL) using a 60x/1.4 Apo oil objective (Olympus). Images were acquired using the ‘optical axis integration’ modus of the softWoRx software over a range of 4 µm, which creates the equivalent of a sum projection of the imaged volume. To measure the time difference between the separation of two chromosomes, images were acquired every 3.5 or 7 seconds for 1 hour. To visualize securin-GFP dynamics, images were acquired every 15 seconds for 1.5–2 hours.

### Data processing and analysis

Images were deconvolved using softWoRx software when necessary to improve signal clarity. The time point of sister chromatid separation was scored manually and was defined as the last time point at which sister chromatids moved coordinately or convergently before moving consistently towards opposing spindle poles. Kymographs were assembled using a custom MATLAB script and the contrast was enhanced for easier visualization of the separation events. ImageJ Trackmate with manual corrections was used for tracking the kinetics of sister chromatid separation (Ershov et al., 2022; Tinevez et al., 2017), and custom MATLAB and R scripts were used to further process the data.

Time-series trajectories for tracked sister chromatid separation distances were generated in R. Distances were summarized at each time point by the cell population mean and standard deviation. To reduce frame-to-frame noise, mean trajectories were smoothed using generalized additive models with cubic regression splines. For plots where all 3 chromosomes are shown together, chromosome II traces were merged from both strains, using inverse-variance weighting when per-timepoint standard deviations were defined, and otherwise by averaging the per-strain means.

To determine the kinetics of securin-GFP degradation, fluorescent intensities were quantified from time-lapse images. The nuclear signal of TetR-tdTomato was used to define the nucleus as a region of interest (ROI), and the average GFP signal intensity within the ROI was determined for each time point. The extracellular background was determined by averaging the signal intensity of three ROIs placed outside of the cell and was subtracted from the GFP signal. When more than one ROI was present (e.g. after nuclear division), their average was calculated before subtraction. For quantitative feature extraction from the degradation curves, local slopes were calculated by taking a derivative over 7 consecutive points of a smoothed spline (the time point ± 3 frames). The onset of securin degradation was defined as the time point before this local slope repeatedly (>5 frames) dropped below 20 % of a reference slope, which was calculated at 50 % GFP intensity. The curves were normalized by setting the GFP level at degradation onset to 100 % and the minimum of the smoothed curve to 0 %. The normalized degradation rate was then approximated from the linear decay between 60 and 40 % signal. The percentage of securin left at sister chromatid separation, the time between degradation onset and sister chromatid separation, and time points at which 90 % of securin had been degraded were calculated using the normalized curves.

### Statistical analysis

Statistical analysis was performed using R, and results are listed in **Supplementary Table S1**. The Gaussian distribution for each data set was generated using R’s built-in normal density function (dnorm). Histograms were generated using a fixed bin width of 7 seconds, and y-axes were normalized to percent of cells. Datasets were compared by two-sample Kolmogorov-Smirnov test unless stated otherwise.

## Acknowledgments

We thank Saahil Golia and Tatiana Boluarte for help with strain construction and image analysis; Douglas Weidemann for scripts; Takeshi Sakuno, Yoshinori Watanabe, and Yasushi Hiraoka for strains; Sarah Gilmour and SaraH Zanders for long-read sequencing and ddPCR of centromere-tagged strains; as well as the Boehringer Ingelheim Fonds (fellowship to J.K.). Research reported in this publication was supported by the National Institute of General Medical Sciences of the National Institutes of Health under award numbers R35GM119723 (S.H.), R35GM149565 (S.H.), and R35GM138370 (J.C.).

## Author contributions

Conceptualization, J.K. and S.H.; Software, W.W., K.P., J.C., and S.L.; Formal Analysis, W.W., K.P., J.C., S.L., J.K., and S.H.; Investigation, W.W., J.K., and S.H.; Writing – Original Draft, W.W., J.K., and S.H.; Writing – Review & Editing, K.P., J.C., and S.L.; Visualization, W.W. and J.K.; Supervision, J.C. and S.H.; Funding Acquisition, J.C. and S.H.

## Competing Interest

None declared

## Notes

### Competing Interest Statement

The authors have declared no competing interest.

