## Supplemental Text, Tables, Figures for "Chromosome segregation synchrony in *S. pombe* is noise-limited and arises without positive feedback"

#### Deterministic models for separase release

##### 1. Model with positive feedback on securin degradation

*Model description*: The model with feedback on securin degradation is depicted in Fig. 3D. Securin (Sec) and separase (Sep) reversibly form a complex (SecSep). Upon initiation of anaphase, the anaphase-promoting complex (APC/C) activity increases from 0 to a positive value, and securin is regulated by APC/C-mediated degradation. The APC/C activity is assumed to be zero in the basal state ($k_{APC/C}$ = 0), and the SecSep complex is assumed to be in equilibrium with the free proteins. Securin is assumed to be in excess over separase prior to anaphase, and the amount of free separase is assumed to be negligible, i.e., the total concentrations of securin ($Sec_{tot}$) and separase ($Sep_{tot}$) are assumed to be much larger than the dissociation constant of the complex ($K_D = k_{off} / k_{on}$). Hence, the initial equilibrium can be approximated as

$$\begin{aligned} Sec &\approx Sec_{tot} - Sep_{tot} \\ Sep &\approx 0 \\ SecSep &\approx Sep_{tot} \end{aligned} \tag{1}$$

Separase, once released from securin, accelerates the degradation of securin. The differential equations describing this scenario are given by

$$\begin{aligned} \frac{dSec}{dt} &= -k_{on} \cdot Sec \cdot Sep + k_{off} \cdot SecSep - k_{APC/C} \cdot Sec \cdot \left( k_{basal} + k_{FB} \frac{Sep^h}{Sep^h + {K_{FB,50}}^h} \right) \\ \frac{dSep}{dt} &= -k_{on} \cdot Sec \cdot Sep + k_{off} \cdot SecSep + k_{APC/C} \cdot SecSep \cdot \left( k_{basal} + k_{FB} \frac{Sep^h}{Sep^h + {K_{FB,50}}^h} \right) \\ \frac{dSecSep}{dt} &= k_{on} \cdot Sec \cdot Sep - k_{off} \cdot SecSep - k_{APC/C} \cdot SecSep \cdot \left( k_{basal} + k_{FB} \frac{Sep^h}{Sep^h + {K_{FB,50}}^h} \right) \end{aligned} \tag{2}$$

The positive feedback (FB) is assumed to be non-linear and is described using the Hill equation.

*Numerical simulations*: Numerical simulations were performed using the ODE system in Eq. 2. The initial steady state was calculated by setting $k_{APC/C}$ = 0.  Anaphase was simulated by assuming a sigmoidal increase in APC/C activity

$$k_{APC/C}(t) \approx k_{APC/C,max} \cdot \frac{t^n}{t^n + {t_{50}}^n} \tag{3}$$

The following parameter values were assumed in Fig. 3D: total separase concentration $Sep_{tot}$ = 0.05 μM; total securin concentration $Sec_{tot}$ = 0.1 μM; $k_{APC/C,max}$ = 0.02 $sec^{-1}$; $k_{on}$ = 1 $μM^{-1}$ $sec^{-1}$; $k_{off}$ = $10^{-4}$ $sec^{-1}$; $t_{50}$ = 100 sec; n = 10; $k_{basal}$ = 1; $k_{FB}$ = 10 (dotted lines) or $k_{FB}$ = 0 (solid lines); $K_{FB,50}$ = 0.02 μM; h = 2.

#### 2. Model with separase auto-activation

*Model description*: The model with feedback amplification of separase activity is depicted in Fig. 4B. Free separase (Sep), once released from securin (Sec), performs auto-cleavage to generate a highly active separase species ($Sep_{active}$), which is more efficient in auto- and cohesin cleavage. The differential equations describing this scenario are given by

$$\frac{dSec}{dt} = -k_{on} \cdot Sec \cdot Sep + k_{off} \cdot SecSep - k_{APC/C} \cdot Sec$$

$$\frac{dSep}{dt} = -k_{on} \cdot Sec \cdot Sep + k_{off} \cdot SecSep + k_{APC/C} \cdot SecSep - \left(k_{a,1} \cdot Sep^h + k_{a,2} \cdot {Sep_{active}}^h\right) \cdot Sep \quad (4)$$

$$\frac{dSecSep}{dt} = k_{on} \cdot Sec \cdot Sep - k_{off} \cdot SecSep - k_{APC/C} \cdot SecSep$$

$$\frac{dSep_{active}}{dt} = \left(k_{a,1} \cdot Sep^h + k_{a,2} \cdot {Sep_{active}}^h\right) \cdot Sep$$

Auto-cleavage occurs with an exponent h that prevents premature amplification and may arise from oligomerization of separase.

The APC/C activity ($k_{APC/C}$) and the irreversible cleavage reactions ($k_{a,1}$, $k_{a,2}$) are assumed to be zero in the basal state. The basal concentrations of securin, separase and of the SecSep complex are the same as in Eq. 1.

Anaphase was simulated by assuming a sigmoidal increase in APC/C activity (Eq. 3). The following parameter values were assumed in Fig. 4B and S6A: total separase concentration $Sep_{tot}$ = 0.05 μM; total securin concentration $Sec_{tot}$ = 0.1 μM; $k_{APC/C,max}$ = 0.02 $sec^{-1}$ (WT) or 0.0066 $sec^{-1}$ (APC/C mut); $k_{on}$ = $10^{-3}$ $μM^{-1}$ $sec^{-1}$; $k_{off}$ = $10^{-7}$ $sec^{-1}$; $t_{50}$ = 100 sec; n = 10; $k_{a,1}$ = 0.1; $k_{a,2}$ = 300; h = 2.

#### Stochastic models for separase-mediated cohesin cleavage

##### 1. Stochastic models for cohesin cleavage

To understand the experimentally observed distributions of chromosome separation times, we modeled the stochastic cleavage of cohesin complexes that hold sister chromatids together. The model assumes that the *i*-th chromosome begins with an initial number of cohesin complexes, $N_i$, and its sister chromatids separate once its cohesin count decays to a threshold $n_i$. Cohesin cleavage is modeled as an inhomogeneous Poisson process. The rate constant of cohesin cleavage is assumed to be identical across all chromosomes and to increase gradually over a timeframe $\tau$, reflecting the gradual release and/or activation of separase in the cell (Eq. 5).

$$k(t) = \begin{cases} k_{max}\dfrac{t}{\tau}, & 0 \le t \le \tau, \\ k_{max}, & t > \tau. \end{cases} \tag{5}$$

The primary output of the model is the distribution of two pairwise separation-time differences, $T_{12} = T_1 - T_2$ and $T_{32} = T_3 - T_2$ where $T_i$ denotes the separation time for chromosome *i*.

The perturbed cases are assumed to undergo the following parameter changes:

***MBC treatment***: Destabilization of microtubules by MBC is expected to reduce the pulling force on sister chromatids and thereby cause a decrease in the threshold number of cohesins that are required to hold sister chromatids together. Since this effect applies to all three chromosomes, a common multiplicative factor, $\alpha < 1$, is applied to all three threshold values, $\{n_1, n_2, n_3\}$.

***Separase mutation****:* Mutated separase is expected to have lower catalytic activity, corresponding to a lower $k_{max}$. To represent this effect, a multiplicative factor, $\beta_k < 1$, is applied to $k_{max}$.

***APC/C mutation****:* Mutated APC/C is expected to degrade securin more slowly and elongate the ramp phase for separase activity. To represent this effect, a multiplicative factor, $\beta_\tau > 1$, is applied to $\tau$.

***Velcade treatment****:* Velcade has a similar effect to the APC/C mutation. To represent this effect, a multiplicative factor, $\beta_{\tau 2} > 1$, is applied to $\tau$.

Additionally, we implemented two mechanistic variants that encode different hypotheses about cohesin cleavage:

***Processive separase action***: This model variant assumes processive cohesin cleavage with a fixed burst size $b$ (number of cohesins cleaved and removed within one event).

***Steric hindrance*****:** This model variant considers steric hindrance by packed chromosomes that lowers accessibility of separase molecules to cohesins in the interior of chromosomes. For simplicity, we assume that cohesin complexes are distributed in a sphere and that only those at the surface are accessible. As outer cohesin complexes get cleaved, inner complexes are exposed. Under these assumptions, the effective cleavage rate scales with the surface-to-volume ratio of the region occupied by the remaining cohesin. Assuming a uniform density of cohesin in the sphere, the volume of the sphere scales with the number of remaining cohesin complexes, i.e., $V \sim N$. Consequently, the surface-to-volume ratio scales as $\frac{A}{V} \sim V^{-\frac{1}{3}} \sim N^{-\frac{1}{3}}$. Therefore, in this model variant, we assume (Eq. 6)

$$k_{eff}(N,t) = \begin{cases} k(t), & N \leq n_{inner}, \\ k(t)\left(\dfrac{N}{n_{inner}}\right)^{-1/3}, & N > n_{inner}, \end{cases} \tag{6}$$

where $n_{inner}$ is the number of cohesins accommodated by the innermost core, a region with no more steric hindrance effect.

Additionally, to evaluate if positive feedback causes a sharp increase in separase activity (Fig. 4B), we further considered a variation of the three models above, in which the time duration for the rate increase, $\tau$, is constrained at low values ($\tau < 5\ sec$).

The six models resulting from the combinations above are summarized in **Supplementary Table S3**. They were each fitted to the experimental data (wild type and perturbations). The bounds for parameter fitting are listed in **Supplementary Table S4**.

**2. Stochastic simulation algorithms**

**2.1 Gillespie stochastic simulation**

To simulate cohesin cleavage, we implemented a modified Gillespie algorithm. Each simulation tracks the number of cohesin complexes on each of the chromosomes and records the time when each chromosome reaches its corresponding cohesin count threshold for separation.

To accommodate the time-dependent degradation rate constant (Eq. 5), we modified the formula for sampling the next-reaction time from the classical Gillespie formula, $\Delta T = T_{next} - T_{prev} = -\frac{\ln(r)}{k_{max}N_{total}}$, to (Eq. 7)

$$\Delta T = T_{next} - T_{prev} = \sqrt{-\frac{2\tau \cdot \ln(r)}{k_{max}N_{total}} + T_{prev}^2} - T_{prev}, \tag{7}$$

where $T_{next}$ is the next degradation time to be sampled, $T_{prev}$ is the previous degradation time, $N_{total}$ is the sum of the number of cohesin complexes currently remaining on the three chromosomes, and $r$ is a uniform random number between 0 and 1. After sampling for the next cleavage time, the cohesin cleavage is randomly chosen to happen to the *i*-th chromosome with probability $N_i/N_{total}$. For the processive separase action model, every cleavage event removes $b$ cohesin complexes on the chosen chromosome. For the steric hindrance model, $N_{total}$ is modified to the sum of the currently accessible number of cohesin complexes on all chromosomes, where the accessible number is, $N_{i,access} = N_i \cdot (N_i/n_{inner})^{-1/3}$ for $N_i > n_{inner}$ and $N_{i,access} = N_i$ for $N_i \leq n_{inner}$; chromosome selection probability is also modified to $N_{i,access}/N_{total}$.

Note that Eq. 7 gives the inverse-CDF sampling for the next-reaction time with CDF, $P(\Delta T < t) = 1 - \exp(-k_{max}N_{total}\left[(T_{prev} + t)^2 - T_{prev}^2\right]/2\tau)$, which can be derived from Eq. 5 through $P(\Delta T < t) = 1 - \exp\left(-\int_{T_{prev}}^{T_{prev}+t} N_{total}k(t')dt'\right)$. Beyond the ramp phase, the regular Gillespie algorithm (Gillespie, 1976, 1977) based on rate constant $k_{max}$ was used.

The Gillespie simulation was used to generate sample time trajectories (Fig. 5F) and to benchmark the fast simulation method described below.

##### 2.2 Fast simulation

To enable efficient parameter optimization and large-scale validation, we simulated the cohesin degradation dynamics in three chromosomes separately (based on their mutual independence) and adopted alternative exact simulation methods, using either order statistics (models without steric hindrance) or vectorized sum of waiting times approach (model with steric hindrance). These methods are statistically exact, alike the Gillespie algorithm, but offer considerable computational speedups (100x-1000x), allowing scalable model fitting.

*Accelerated sampling via order statistics:* For the basic model, the cleavage of individual cohesins are independent events following an identical waiting time distribution with the following CDF,

$$P(T_{wait} < t) = \begin{cases} 1 - \exp\left(-\frac{k_{max}t^2}{2\tau}\right), & 0 \leq t \leq \tau, \\ 1 - \exp\left(-\frac{k_{max}\tau}{2} - k_{max}(t-\tau)\right), & t > \tau. \end{cases} \quad (8)$$

The separation time for the *i*-th chromosome is the $(n_i + 1)$-th largest cohesin cleavage time out of a total of $N_i$ cleavage times that follow the distribution in Eq. 8. As each cleavage time can be converted from a uniform random number through CDF inverse, which is a monotonically increasing function, the $(n_i + 1)$-th largest cleavage time is the CDF inverse of the $(n_i + 1)$-th largest value of $N_i$ uniform random numbers. The latter is known to follow the beta distribution $Beta(N_i - n_i, n_i + 1)$. Therefore, by directly drawing one random number from this beta distribution and converting it through the inverse of Eq. 8, we obtain a sample separation time of a chromosome with a highly efficient $O(1)$ operation.

The accelerated sampling method applies to the processive separase action model, too, when cohesin complexes are treated as predefined groups. A single cleavage event eliminates one group of $b$ cohesins. To utilize the order statistics framework, we transform the parameters as effective counts:

$$\hat{N}_i = \left\lceil \frac{N_i}{b} \right\rceil, \hat{n}_i = \hat{N}_i - \left\lceil \frac{N_i - n_i}{b} \right\rceil. \quad (9)$$

The process then proceeds as the non-processive case with $\hat{N}_i$ and $\hat{n}_i$.

*Vectorized simulation:* For the steric hindrance model, the effective rate constant depends on the system's current state (Eq. 6). Therefore, cleavage events of individual cohesin complexes are no longer independent of each other, making the fast-sampling method invalid. To accelerate the simulation, we take advantage of the convenient feature that there is only one reaction in the system and employ vectorization across simulations. Instead of simulating one trajectory at a time, we simultaneously simulate $M$ independent cohesin loss trajectories. For each step from state $N$ to $N-1$, we use Eq. 7 to determine the next event times across all $M$ simulations. This approach leverages efficient array operations to achieve a significant speedup.

**3. Parameter estimation and model selection**

*Objective metric — Earth Mover's Distance (EMD).*

For each candidate parameter set, the simulation methods described above generated a batch of $M$ = 10,000 independent samples of $T_{12}$ and $T_{32}$, respectively, for each experimental scenario. Model fit was assessed using the Earth Mover's Distance (EMD; Wasserstein-1 distance) (Bazán et al., 2019; Rubner et al., 1998), a robust goodness-of-fit metric that quantifies the similarity between the empirical distribution of experimental data and that of the simulated data.

The objective function was defined as the summed EMD values across five experiments (control, MBC, separase mutant, APC/C mutant, velcade), which is also termed aggregate EMD. Sample size of 10,000 was confirmed to generate a sufficiently precise estimate of EMD (Fig. S8A).

*Optimizer — Differential evolution (global) + local refinement.*

Model parameters were estimated using a two-stage optimization protocol. First, a global search was performed using differential evolution, an efficient population-based stochastic algorithm that explores the parameter space (Storn and Price, 1997). The algorithm was implemented using the `scipy.optimize.differential_evolution()` function in the SciPy library. We used the 'best1bin' strategy with a population size of 10 (corresponding to 10x the number of parameters in each model variant), mutation factor (0.5, 1.0), recombination constant 0.7, and a relative convergence tolerance of 0.01. The top-performing candidate solutions were then refined locally using the L-BFGS-B algorithm (Byrd et al., 1995) to maximize fit quality. Choices of population size and convergence tolerance were informed by benchmark results shown in Fig. S8B,C.

*Cross-validation.*

To reduce bias and avoid overfitting, 5-fold cross-validation was performed on the experimental dataset. Data were split into five stratified folds; for each fold, optimization was conducted on 80 % of the data, and EMD was computed on the held-out 20 %. Model comparison between mechanisms in Fig. 5C was performed using the aggregate cross-validated EMD metric (mean ± SE across folds).

##### 4. OAT parameter sensitivity analysis

To evaluate the impact of each parameter on separation synchrony, we performed a One-At-a-Time (OAT) sensitivity analysis. Each parameter was perturbed across the range of this parameter while holding all others fixed at the previously determined optimal value. For each parameter set we ran 10 batches of 10,000 separation time difference ($\Delta t$) simulations.

##### 5. Algorithm implementation

All simulations and optimization procedures were implemented in Python using standard scientific libraries. Computationally intensive parameter fitting and cross-validation were performed in parallel on a high-performance computing cluster. Parameter estimates are reported from best-fit solutions with uncertainties derived from cross-validated ensembles and model performance was summarized by the mean cross-validated aggregate EMD. All analyses are reproducible using the archived code and parameter settings.

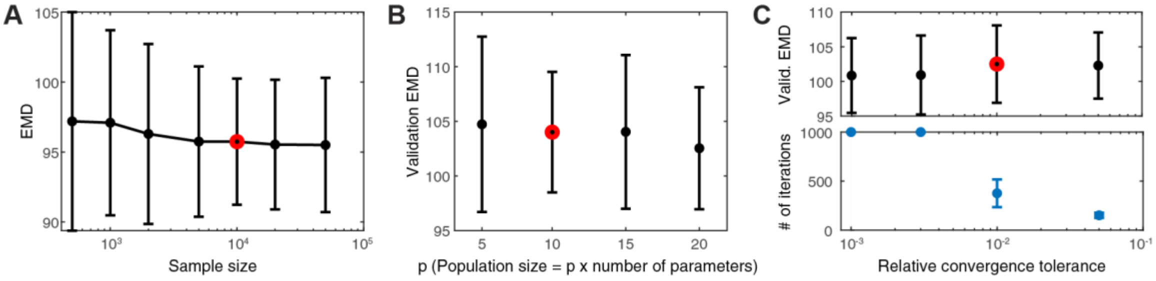

**Figure S8. Benchmarks and selection of key stochastic simulation and optimization hyperparameters.** **(A)** EMD estimate is unbiased, exhibiting a stable mean across sample sizes, and its precision (standard error) stabilizes beyond a sample size of 10,000. The optimal parameter set fitted to the complete experimental dataset was used, yielding lower EMD values than the five-fold cross-validation EMDs shown in (B) and (C). **(B)** Varying the population size in differential evolution has no significant effect on the optimization results. We therefore used p = 10 in this study. Five-fold cross-validation EMD values are shown. **(C)** Mean and standard error of five-fold cross-validation EMD are stable across the tested range of relative tolerances (upper panel). However, because stochastic simulations are limited by finite sampling, convergence is constrained by simulation sample size. Consequently, stringent relative tolerances (< 0.01) trigger spurious non-convergence warnings and incur unnecessary computational cost (lower panel; maximum iteration limit reached for the two smallest tolerances) after the practical convergence limit has already been achieved. We therefore selected a relative tolerance of 0.01. Error bars represent mean ± SE. Red circles indicate the hyperparameter values chosen in this study.

#### Supplementary Table S1 - Statistical analysis of experimental results

| | | n | mean (sec) | median (sec) | stdev (sec) | p-value (Kolmogorov-Smirnov) | tested against |
|---|---|---|---|---|---|---|---|
| **WT (rich medium)** | chromosome I vs. II | 132 | -6.77 | -8 | 16.41 | | |
| | chromosome III vs. II | 86 | 11.5 | 13 | 16.89 | | |
| **WT (minimal medium)** | chromosome I vs. II | 126 | -3.44 | -1.01 | 19 | 0.164 | chromosome I vs. II WT (minimal medium) |
| | chromosome III vs. II | 145 | 11.47 | 12.99 | 13.51 | [1]0.389<br>[2]1.56e-14 | [1]chromosome II vs. III WT (rich medium)<br>[2]chromosome I vs. II WT (rich medium) |
| **separase mutant (*cut1-206,* rich medium)** | chromosome I vs. II | 87 | -7.46 | -8 | 40.55 | 0.048441 | chromosome I vs. II WT (rich medium) |
| | chromosome III vs. II | 71 | 32.69 | 34 | 31.89 | 7.65E-09 | chromosome II vs. III WT (rich medium) |
| **separase mutant (*cut1-206,* rich to minimal medium)** | chromosome I vs. II | 74 | -4.4 | -3.5 | 28.52 | [1]0.036064<br>[2]0.7575 | [1]chromosome I vs. II WT (rich medium)<br>[2]chromosome I vs. II separase mutant (rich medium) |
| | chromosome III vs. II | 188 | 24.61 | 24.5 | 34.51 | [1]2.45e−9<br>[2]0.1219 | [1]chromosome II vs. III WT (rich medium)<br>[2]chromosome II vs. III separase mutant (rich medium) |
| **separase mutant (*cut1-206,* combined results)** | chromosome I vs. II | 161 | -6.05 | -8 | 35.46 | [1]0.01852<br>[2]0.9993<br>[3]0.9962 | [1]chromosome I vs. II WT (rich medium)<br>[2]chromosome I vs. II separase mutant (rich medium)<br>[3]chromosome I vs. II separase mutant (rich to minimal medium) |
| | chromosome III vs. II | 259 | 26.83 | 28 | 33.94 | [1]2.94e-11<br>[2]0.4016<br>[3]0.979857 | [1]chromosome II vs. III WT (rich medium)<br>[2]chromosome II vs. III separase mutant (rich medium)<br>[3]chromosome II vs. III separase mutant (rich to minimal medium) |
| **securin overexpression** | chromosome I vs. II | 62 | -11.29 | -14 | 21.58 | 0.002129 | chromosome I vs. II WT (minimal medium) |
| | chromosome III vs. II | 95 | 9.95 | 10.5 | 13 | 0.378779 | chromosome II vs. III WT (minimal medium) |
| **securin/separase overexpression** | chromosome I vs. II | 99 | -8.63 | -7 | 19.56 | [1]0.01304<br>[2]0.023857 | [1]chromosome I vs. II securin overexpression<br>[2]chromosome I vs. II WT (minimal medium) |
| | chromosome III vs. II | 60 | 7.53 | 3.5 | 18.81 | [1]0.124161<br>[2]0.002022 | [1]chromosome II vs. III securin overexpression<br>[2]chromosome II vs. III WT (minimal medium) |
| **WT + 3 ug/mL MBC (minimal medium)** | chromosome I vs. II | 67 | -11.54 | -7 | 21.78 | 0.00066 | chromosome I vs. II WT (minimal medium) |
| | chromosome III vs. II | 62 | 14.79 | 10.5 | 22.1 | 0.0896 | chromosome II vs. III WT (minimal medium) |
| **klp5Δ** | chromosome I vs. II | 56 | -3.69 | -1.75 | 18.63 | 0.803657 | chromosome I vs. II WT (minimal medium) |
| | chromosome III vs. II | 37 | 13.15 | 10.5 | 14.3 | 0.332162 | chromosome II vs. III WT (minimal medium) |
| **APC/C mutant (*cut9-665*, rich medium)** | chromosome I vs. II | 158 | -11.9 | -12.25 | 43.51 | 0.004576 | chromosome I vs. II WT (rich medium) |
| | chromosome III vs. II | 161 | 18.79 | 17.5 | 24.99 | 0.00023 | chromosome II vs. III WT (rich medium) |

| | | | | | | | |
|---|---|---|---|---|---|---|---|
| **proteasome inhibition (minimal medium)** | chromosome I vs. II | 131 | -9.62 | -14 | 91.8 | 7.58E-07 | chromosome I vs. II WT (minimal medium) |
| | chromosome III vs. II | 153 | 41.07 | 35 | 52.67 | 4.24E-20 | chromosome II vs. III WT (minimal medium) |
| **ΔN-cyclin B normal (minimal medium)** | chromosome I vs. II | 101 | -5.1 | -7 | 18.12 | 0.263528 | chromosome I vs. II WT (minimal medium) |
| **ΔN-cyclin B pseudo-meta (minimal medium)** | chromosome I vs. II | 98 | -5.95 | -8 | 24.6 | [1]0.237701<br>[2]0.824825 | [1]chromosome I vs. II WT (minimal medium)<br>[2]ΔN-cyclin B normal |
| **imr1L / dh1R** | imr1L vs dh1R | 78 | 0.2 | 0 | 7.3 | | |
| | imr1L vs dh1R | 88 | 0.9 | 0 | 4.1 | | |
| **WT central centromere tags** | chromosome I vs. II | 137 | -3.49 | -3.5 | 17.41 | 0.4819 | chromosome I vs II WT (minimal medium) |
| | chromosome III vs. II | 146 | 7.77 | 7.12 | 18.87 | 0.0291 | chromosome II vs III WT (minimal medium) |
| **marker swap** | chromosome I vs. II | 152 | -8.96 | -7 | 19.16 | 1.54E-04 | chromosome I vs II WT (minimal medium) |
| **arm markers** | chromosome II vs. arm | 79 | 120.75 | 119 | 46.07 | [1]<2.23e-308<br>[2]<2.23e-308<br>[3]0.1141 | [1]chromosome I vs II WT (minimal medium)<br>[2]chromosome II vs III WT (minimal medium)<br>[3]central centromere II vs arm |
| | central centromere II vs. arm | 72 | 105.17 | 103.75 | 46.63 | [1]<2.23e-308<br>[2]<2.23e-308 | [1]chromosome I vs II WT (minimal medium)<br>[2]chromosome II vs III WT (minimal medium) |

### Supplementary Table S2 - *S. pombe* strains

**Figure 1C-D**

| | | |
|---|---|---|
| SL239' | *h+* | *leu1- ade6-M216 cen2::lacO-kanR:ura+ his7+::P.dis1-GFP-lacI-NLS cen3::Scer\LEU2:tetO zfs1+::natR:P.adh31-tetR-tdTomato* |

**Figure 1E**

| | | |
|---|---|---|
| SU495/SU496 | *h-* | *leu1- ura4-D18? FY397::RS imr1L::lacO-RS his7+::P.dis1-GFP-lacI-NLS dh1R::tetO:ura4+ zfs1+::natR:P.adh31-tetR-tdTomato* |

**Figure 1F**

| | | |
|---|---|---|
| SI541 | *h+* | *leu1- ade6-M216 ura4-D18? cen2::lacO-kanR:ura+ his7+::P.dis1-GFP-lacI-NLS dh1L::ura4+:tetO zfs1+::natR:P.adh31-tetR-tdTomato* |
| SL239' | *h+* | *leu1- ade6-M216 cen2::lacO-kanR:ura+ his7+::P.dis1-GFP-lacI-NLS cen3::Scer\LEU2:tetO zfs1+::natR:P.adh31-tetR-tdTomato* |

**Figure 1H**

| | | |
|---|---|---|
| SI541 | *h+* | *leu1- ade6-M216 ura4-D18? cen2::lacO-kanR:ura+ his7+::P.dis1-GFP-lacI-NLS dh1L::ura4+:tetO zfs1+::natR:P.adh31-tetR-tdTomato* |
| SL239' | *h+* | *leu1- ade6-M216 cen2::lacO-kanR:ura+ his7+::P.dis1-GFP-lacI-NLS cen3::Scer\LEU2:tetO zfs1+::natR:P.adh31-tetR-tdTomato* |
| PQ768/PQ768' | *h+* | *leu1- ura4-D18 cnt1::lacO his7+::P.dis1-GFP-lacI-NLS cnt2::tetO:ura4+ zfs1+::natR:P.adh31-tetR-tdTomato* |
| ST968/ST968' | *h+* | *leu1- ura4-D18 cnt3::Scer\LEU2:lacO his7+::P.dis1-GFP-lacI-NLS cnt2::tetO:ura4+ zfs1+::natR:P.adh31-tetR-tdTomato* |

**Figure 2A-B**

| | | |
|---|---|---|
| SI541 | *h+* | *leu1- ade6-M216 ura4-D18? cen2::lacO-kanR:ura+ his7+::P.dis1-GFP-lacI-NLS dh1L::ura4+:tetO zfs1+::natR:P.adh31-tetR-tdTomato* |
| SL239' | *h+* | *leu1- ade6-M216 cen2::lacO-kanR:ura+ his7+::P.dis1-GFP-lacI-NLS cen3::Scer\LEU2:tetO zfs1+::natR:P.adh31-tetR-tdTomato* |
| SM388 | *h?* | *leu1- ade6-M21? ura4-D18? cen2::lacO-kanR:ura+ his7+::P.dis1-GFP-lacI-NLS dh1L::ura4+:tetO zfs1+::natR:P.adh31-tetR-tdTomato cut1-206* |
| SM387 | *h?* | *leu1- ade6-M21? cen2::lacO-kanR:ura+ his7+::P.dis1-GFP-lacI-NLS cen3::Scer\LEU2:tetO zfs1+::natR:P.adh31-tetR-tdTomato cut1-206* |

**Figure 2C-D**

| | | |
|---|---|---|
| SI541 | *h+* | *leu1- ade6-M216 ura4-D18? cen2::lacO-kanR:ura+ his7+::P.dis1-GFP-lacI-NLS dh1L::ura4+:tetO zfs1+::natR:P.adh31-tetR-tdTomato* |
| SL239' | *h+* | *leu1- ade6-M216 cen2::lacO-kanR:ura+ his7+::P.dis1-GFP-lacI-NLS cen3::Scer\LEU2:tetO zfs1+::natR:P.adh31-tetR-tdTomato* |
| SW562 | *h-* | *leu1+::P.cut1(long)-cut1+ cen2::lacO-kanR:ura+ his7+::P.dis1-GFP-lacI-NLS dh1L::ura4+:tetO zfs1+::natR:P.adh31-tetR-tdTomato natNT2::P.adh1(#6)-cut2+-GFP(Y66L):kanR* |
| SW566 | *h-* | *leu1+::P.cut1(long)-cut1+ cen2::lacO-kanR:ura+ his7+::P.dis1-GFP-lacI-NLS cen3::Scer\LEU2:tetO zfs1+::natR:P.adh31-tetR-tdTomato natNT2::P.adh1(#6)-cut2+-GFP(Y66L):kanR* |

**Figure2E-F**

| | | |
|---|---|---|
| SI541 | *h+* | *leu1- ade6-M216 ura4-D18? cen2::lacO-kanR:ura+ his7+::P.dis1-GFP-lacI-NLS dh1L::ura4+:tetO zfs1+::natR:P.adh31-tetR-tdTomato* |
| SL239' | *h+* | *leu1- ade6-M216 cen2::lacO-kanR:ura+ his7+::P.dis1-GFP-lacI-NLS cen3::Scer\LEU2:tetO zfs1+::natR:P.adh31-tetR-tdTomato* |

**Figure 3A-B**

| | | |
|---|---|---|
| SL274 | *h+* | *leu1+::P.nmt81-cdc13Δ(1-67) ade6-M216 ura4-D18? cen2::lacO-kanR:ura+ his7+::P.dis1-GFP-lacI-NLS dh1L::ura4+:tetO zfs1+::natR:P.adh31-tetR-tdTomato* |
| SI541 | *h+* | *leu1- ade6-M216 ura4-D18? cen2::lacO-kanR:ura+ his7+::P.dis1-GFP-lacI-NLS dh1L::ura4+:tetO zfs1+::natR:P.adh31-tetR-tdTomato* |

**Figure 3E-F**

| | | |
|---|---|---|
| SL249 | *h-* | *leu1- ade6-M216 dh1L::ura4+:tetO zfs1+::natR:P.adh31-tetR-tdTomato cut2+-GFP:kanR* |
| SL275 | *h-* | *leu1+::P.nmt81-cut2Δ(1-80) ade6-M216 dh1L::ura4+:tetO zfs1+::natR:P.adh31-tetR-tdTomato cut2+-GFP:kanR* |

**Figure 4C**

| | | |
|---|---|---|
| SI541 | *h+* | *leu1- ade6-M216 ura4-D18? cen2::lacO-kanR:ura+ his7+::P.dis1-GFP-lacI-NLS dh1L::ura4+:tetO zfs1+::natR:P.adh31-tetR-tdTomato* |
| SL239' | *h+* | *leu1- ade6-M216 cen2::lacO-kanR:ura+ his7+::P.dis1-GFP-lacI-NLS cen3::Scer\LEU2:tetO zfs1+::natR:P.adh31-tetR-tdTomato* |
| SL231 | *h+* | *leu1- ade6-M216 cen2::lacO-kanR:ura+ his7+::P.dis1-GFP-lacI-NLS dh1L::ura4+:tetO zfs1+::natR:P.adh31-tetR-tdTomato cut9-665* |
| SX444 | *h?* | *leu1- ura4-D18? cen2::lacO-kanR:ura+ his7+::P.dis1-GFP-lacI-NLS dh1L::ura4+:tetO zfs1+::natR:P.adh31-tetR-tdTomato cut9-665* |
| SX445' | *h?* | *leu1- ade6-M216 ura4-D18? cen2::lacO-kanR:ura+ his7+::P.dis1-GFP-lacI-NLS dh1L::ura4+:tetO zfs1+::natR:P.adh31-tetR-tdTomato cut9-665* |
| SM386/SM386'/ SM386'' | *h?* | *leu1- ade6-M216 cen2::lacO-kanR:ura+ his7+::P.dis1-GFP-lacI-NLS cen3::Scer\LEU2:tetO zfs1+::natR:P.adh31-tetR-tdTomato cut9-665* |
| **Figure 4D** | | |
| SI541 | *h+* | *leu1- ade6-M216 ura4-D18? cen2::lacO-kanR:ura+ his7+::P.dis1-GFP-lacI-NLS dh1L::ura4+:tetO zfs1+::natR:P.adh31-tetR-tdTomato* |
| SL239' | *h+* | *leu1- ade6-M216 cen2::lacO-kanR:ura+ his7+::P.dis1-GFP-lacI-NLS cen3::Scer\LEU2:tetO zfs1+::natR:P.adh31-tetR-tdTomato* |
| **Figure 5B** | | |
| SI541 | *h+* | *leu1- ade6-M216 ura4-D18? cen2::lacO-kanR:ura+ his7+::P.dis1-GFP-lacI-NLS dh1L::ura4+:tetO zfs1+::natR:P.adh31-tetR-tdTomato* |
| SL239' | *h+* | *leu1- ade6-M216 cen2::lacO-kanR:ura+ his7+::P.dis1-GFP-lacI-NLS cen3::Scer\LEU2:tetO zfs1+::natR:P.adh31-tetR-tdTomato* |
| **Figure S1A-D** | | |
| SL249 | *h-* | *leu1- ade6-M216 dh1L::ura4+:tetO zfs1+::natR:P.adh31-tetR-tdTomato cut2+-GFP:kanR* |
| **Figure S1E** | | |
| SI541 | *h+* | *leu1- ade6-M216 ura4-D18? cen2::lacO-kanR:ura+ his7+::P.dis1-GFP-lacI-NLS dh1L::ura4+:tetO zfs1+::natR:P.adh31-tetR-tdTomato* |
| **Figure S1F** | | |
| SU495 | *h-* | *leu1- ura4-D18? FY397::RS imr1L::lacO-RS his7+::P.dis1-GFP-lacI-NLS dh1R::tetO:ura4+ zfs1+::natR:P.adh31-tetR-tdTomato* |
| **Figure S1G** | | |
| SI541 | *h+* | *leu1- ade6-M216 ura4-D18? cen2::lacO-kanR:ura+ his7+::P.dis1-GFP-lacI-NLS dh1L::ura4+:tetO zfs1+::natR:P.adh31-tetR-tdTomato* |
| SL239' | *h+* | *leu1- ade6-M216 cen2::lacO-kanR:ura+ his7+::P.dis1-GFP-lacI-NLS cen3::Scer\LEU2:tetO zfs1+::natR:P.adh31-tetR-tdTomato* |
| **Figure S1H** | | |
| SI541 | *h+* | *leu1- ade6-M216 ura4-D18? cen2::lacO-kanR:ura+ his7+::P.dis1-GFP-lacI-NLS dh1L::ura4+:tetO zfs1+::natR:P.adh31-tetR-tdTomato* |
| ST653'' | *h+* | *leu1- ura4-D18? dh1L::lacO:ura4+ his7+::P.dis1-GFP-lacI-NLS cen2::Scer\LEU2:tetO zfs1+::natR:P.adh31-tetR-tdTomato* |
| **Figure S1I,K** | | |
| SU287 | *h90* | *leu1- ura4-D18? ade8∆::kanR:ura4:lacO his7+::P.dis1-GFP-lacI-NLS cnt2::tetO:ura4+ zfs1+::natR:P.adh31-tetR-tdTomato* |
| **Figure S1J** | | |
| SU286 | *h90* | *leu1- ura4-D18? ade8∆::kanR:ura4:lacO his7+::P.dis1-GFP-lacI-NLS cen2::Scer\LEU2:tetO zfs1+::natR:P.adh31-tetR-tdTomato* |
| **Figure S2A-B,E** | | |
| SI541 | *h+* | *leu1- ade6-M216 ura4-D18? cen2::lacO-kanR:ura+ his7+::P.dis1-GFP-lacI-NLS dh1L::ura4+:tetO zfs1+::natR:P.adh31-tetR-tdTomato* |
| SL239' | *h+* | *leu1- ade6-M216 cen2::lacO-kanR:ura+ his7+::P.dis1-GFP-lacI-NLS cen3::Scer\LEU2:tetO zfs1+::natR:P.adh31-tetR-tdTomato* |
| SM388 | *h?* | *leu1- ade6-M21? ura4-D18? cen2::lacO-kanR:ura+ his7+::P.dis1-GFP-lacI-NLS dh1L::ura4+:tetO zfs1+::natR:P.adh31-tetR-tdTomato cut1-206* |
| SM387 | *h?* | *leu1- ade6-M21? cen2::lacO-kanR:ura+ his7+::P.dis1-GFP-lacI-NLS cen3::Scer\LEU2:tetO zfs1+::natR:P.adh31-tetR-tdTomato cut1-206* |

**Figure S2C-D**

| | | |
|---|---|---|
| SM388 | *h?* | *leu1- ade6-M21? ura4-D18? cen2::lacO-kanR:ura+ his7+::P.dis1-GFP-lacI-NLS dh1L::ura4+:tetO zfs1+::natR:P.adh31-tetR-tdTomato cut1-206* |

**Figure S3A-B**

| | | |
|---|---|---|
| SL249 | *h-* | *leu1- ade6-M216 dh1L::ura4+:tetO zfs1+::natR:P.adh31-tetR-tdTomato cut2+-GFP:kanR* |
| SM325' | *h-* | *leu1- ade6-M216 ura4-D18? dh1L::ura4+:tetO zfs1+::natR:P.adh31-tetR-tdTomato natNT2::P.adh1(#6)-cut2-GFP:kanR* |
| SW503 | *h-* | *leu1- ade6-M216 ura4-D18? dh1L::ura4+:tetO zfs1+::natR:P.adh31-tetR-tdTomato natNT2::P.adh1(#6)-cut2-GFP:kanR hphNT1::P.ark1-cut1+* |
| SW502 | *h-* | *leu1+::P.cut1(long)-cut1+ ade6-M216 ura4-D18? dh1L::ura4+:tetO zfs1+::natR:P.adh31-tetR-tdTomato natNT2::P.adh1(#6)-cut2-GFP:kanR* |

**Figure S3C,E**

| | | |
|---|---|---|
| SI541 | *h+* | *leu1- ade6-M216 ura4-D18? cen2::lacO-kanR:ura+ his7+::P.dis1-GFP-lacI-NLS dh1L::ura4+:tetO zfs1+::natR:P.adh31-tetR-tdTomato* |
| SL239' | *h+* | *leu1- ade6-M216 cen2::lacO-kanR:ura+ his7+::P.dis1-GFP-lacI-NLS cen3::Scer\LEU2:tetO zfs1+::natR:P.adh31-tetR-tdTomato* |
| SW560 | *h-* | *leu1- cen2::lacO-kanR:ura+ his7+::P.dis1-GFP-lacI-NLS dh1L::ura4+:tetO zfs1+::natR:P.adh31-tetR-tdTomato natNT2::P.adh1(#6)-cut2+-GFP(Y66L):kanR* |
| SW559 | *h+* | *leu1- cen2::lacO-kanR:ura+ his7+::P.dis1-GFP-lacI-NLS cen3::Scer\LEU2:tetO zfs1+::natR:P.adh31-tetR-tdTomato natNT2::P.adh1(#6)-cut2+-GFP(Y66L):kanR* |

**Figure S3D,F**

| | | |
|---|---|---|
| SW560 | *h-* | *leu1- cen2::lacO-kanR:ura+ his7+::P.dis1-GFP-lacI-NLS dh1L::ura4+:tetO zfs1+::natR:P.adh31-tetR-tdTomato natNT2::P.adh1(#6)-cut2+-GFP(Y66L):kanR* |
| SW559 | *h+* | *leu1- cen2::lacO-kanR:ura+ his7+::P.dis1-GFP-lacI-NLS cen3::Scer\LEU2:tetO zfs1+::natR:P.adh31-tetR-tdTomato natNT2::P.adh1(#6)-cut2+-GFP(Y66L):kanR* |
| SW562 | *h-* | *leu1+::P.cut1(long)-cut1+ cen2::lacO-kanR:ura+ his7+::P.dis1-GFP-lacI-NLS dh1L::ura4+:tetO zfs1+::natR:P.adh31-tetR-tdTomato natNT2::P.adh1(#6)-cut2+-GFP(Y66L):kanR* |
| SW566 | *h-* | *leu1+::P.cut1(long)-cut1+ cen2::lacO-kanR:ura+ his7+::P.dis1-GFP-lacI-NLS cen3::Scer\LEU2:tetO zfs1+::natR:P.adh31-tetR-tdTomato natNT2::P.adh1(#6)-cut2+-GFP(Y66L):kanR* |

**Figure S4A**

| | | |
|---|---|---|
| SW596 | *h-* | *cen3::Scer\LEU2:tetO zfs1+::natR:P.adh31-tetR-tdTomato leu1-32::SV40-GFP-atb2[LEU1]* |

**Figure S4B**

| | | |
|---|---|---|
| SI541 | *h+* | *leu1- ade6-M216 ura4-D18? cen2::lacO-kanR:ura+ his7+::P.dis1-GFP-lacI-NLS dh1L::ura4+:tetO zfs1+::natR:P.adh31-tetR-tdTomato* |
| SL239' | *h+* | *leu1- ade6-M216 cen2::lacO-kanR:ura+ his7+::P.dis1-GFP-lacI-NLS cen3::Scer\LEU2:tetO zfs1+::natR:P.adh31-tetR-tdTomato* |

**Figure S4C**

| | | |
|---|---|---|
| SI541 | *h+* | *leu1- ade6-M216 ura4-D18? cen2::lacO-kanR:ura+ his7+::P.dis1-GFP-lacI-NLS dh1L::ura4+:tetO zfs1+::natR:P.adh31-tetR-tdTomato* |
| SL239' | *h+* | *leu1- ade6-M216 cen2::lacO-kanR:ura+ his7+::P.dis1-GFP-lacI-NLS cen3::Scer\LEU2:tetO zfs1+::natR:P.adh31-tetR-tdTomato* |
| SX614 | *h+* | *leu1- ade6-M216 ura4-D18? cen2::lacO-kanR:ura+ his7+::P.dis1-GFP-lacI-NLS dh1L::ura4+:tetO zfs1+::natR:P.adh31-tetR-tdTomato klp5Δ::hygR* |
| SX615 | *h+* | *leu1- ade6-M216 cen2::lacO-kanR:ura+ his7+::P.dis1-GFP-lacI-NLS cen3::Scer\LEU2:tetO zfs1+::natR:P.adh31-tetR-tdTomato klp5Δ::hygR* |

**Figure S4D**

| | | |
|---|---|---|
| SX614 | *h+* | *leu1- ade6-M216 ura4-D18? cen2::lacO-kanR:ura+ his7+::P.dis1-GFP-lacI-NLS dh1L::ura4+:tetO zfs1+::natR:P.adh31-tetR-tdTomato klp5Δ::hygR* |

**Figure S5A**

| | | |
|---|---|---|
| SL249 | *h-* | *leu1- ade6-M216 dh1L::ura4+:tetO zfs1+::natR:P.adh31-tetR-tdTomato cut2+-GFP:kanR* |
| SL253 | *h-* | *leu1+::P.nmt81-cdc13Δ(1-67) ade6-M216 dh1L::ura4+:tetO zfs1+::natR:P.adh31-tetR-tdTomato cut2+-GFP* |

**Figure S5B**

| | | |
|---|---|---|
| SL249 | *h-* | *leu1- ade6-M216 dh1L::ura4+:tetO zfs1+::natR:P.adh31-tetR-tdTomato cut2+-GFP:kanR* |

| | | |
|---|---|---|
| SL275 | *h-* | *leu1+::P.nmt81-cut2Δ(1-80) ade6-M216 dh1L::ura4+:tetO zfs1+::natR:P.adh31-tetR-tdTomato cut2+-GFP:kanR* |
| **Figure S6B-D** | | |
| SL249 | *h-* | *leu1- ade6-M216 dh1L::ura4+:tetO zfs1+::natR:P.adh31-tetR-tdTomato cut2+-GFP:kanR* |
| SL258' | *h-* | *leu1- ade6-M216 dh1L::ura4+:tetO zfs1+::natR:P.adh31-tetR-tdTomato cut9-665 cut2+-GFP:kanR* |
| **Figure S6E** | | |
| ST968 | *h+* | *leu1- ura4-D18 cnt3::Scer\LEU2:lacO his7+::P.dis1-GFP-lacI-NLS cnt2::tetO:ura4+ zfs1+::natR:P.adh31-tetR-tdTomato* |
| **Figure S7A** | | |
| SI541 | *h+* | *leu1- ade6-M216 ura4-D18? cen2::lacO-kanR:ura+ his7+::P.dis1-GFP-lacI-NLS dh1L::ura4+:tetO zfs1+::natR:P.adh31-tetR-tdTomato* |
| SL239' | *h+* | *leu1- ade6-M216 cen2::lacO-kanR:ura+ his7+::P.dis1-GFP-lacI-NLS cen3::Scer\LEU2:tetO zfs1+::natR:P.adh31-tetR-tdTomato* |
| SM388 | *h?* | *leu1- ade6-M21? ura4-D18? cen2::lacO-kanR:ura+ his7+::P.dis1-GFP-lacI-NLS dh1L::ura4+:tetO zfs1+::natR:P.adh31-tetR-tdTomato cut1-206* |
| SM387 | *h?* | *leu1- ade6-M21? cen2::lacO-kanR:ura+ his7+::P.dis1-GFP-lacI-NLS cen3::Scer\LEU2:tetO zfs1+::natR:P.adh31-tetR-tdTomato cut1-206* |
| SL231 | *h+* | *leu1- ade6-M216 cen2::lacO-kanR:ura+ his7+::P.dis1-GFP-lacI-NLS dh1L::ura4+:tetO zfs1+::natR:P.adh31-tetR-tdTomato cut9-665* |
| SX444 | *h?* | *leu1- ura4-D18? cen2::lacO-kanR:ura+ his7+::P.dis1-GFP-lacI-NLS dh1L::ura4+:tetO zfs1+::natR:P.adh31-tetR-tdTomato cut9-665* |
| SX445' | *h?* | *leu1- ade6-M216 ura4-D18? cen2::lacO-kanR:ura+ his7+::P.dis1-GFP-lacI-NLS dh1L::ura4+:tetO zfs1+::natR:P.adh31-tetR-tdTomato cut9-665* |
| SM386/SM386'/ SM386'' | *h?* | *leu1- ade6-M216 cen2::lacO-kanR:ura+ his7+::P.dis1-GFP-lacI-NLS cen3::Scer\LEU2:tetO zfs1+::natR:P.adh31-tetR-tdTomato cut9-665* |

#### Supplementary Table S3 – Stochastic model variants

| | Features | Parameters | Separase activity ramp |
|---|---|---|---|
| **Basic model** | | N2, n2, R12, R32, r12, r32, $k_{max}$, $\tau$<br><br>Perturbation conditions:<br>$\alpha$, $\beta_k$, $\beta_\tau$, $\beta_{\tau 2}$ | slow, $2 < \tau < 240$<br>fast, $0.5 < \tau < 5$ |
| **Processive separase action** | $b$ cohesin molecules removed within one removal event | same as basic model, and $b$ | slow, $2 < \tau < 240$<br>fast, $0.5 < \tau < 5$ |
| **Steric Hindrance** | effective $k$ scales with surface-to-volume ratio (Eq. 6) | same as basic model, and $n_{inner}$ | slow, $2 < \tau < 240$<br>fast, $0.5 < \tau < 5$ |

#### Supplementary Table S4 - Model parameters and their bounds for fitting

| Symbol | Description | Unit | Parameter Bounds | Parameter Justification |
|---|---|---|---|---|
| $N_2$ | Initial cohesin count for the reference chromosome (chromosome 2) | 1 | [50 - 1000] | Range was set broadly to include (i) cohesin-binding patterns in fission yeast (Schmidt et al., 2009; Mizuguchi et al., 2014), (ii) absolute protein abundance showing cohesin subunits exist at $10^2$-$10^3$ or more molecules per cell (Marguerat et al., 2012; Carpy et al., 2014), and (iii) additional quantification showing large numbers of cohesin complex complexes in other cell types, both cohesive and non-cohesive (Holzmann et al., 2019), and (iv) the fact that not all cohesin contributes to cohesion (Gerlich et al., 2006; Feytout et al., 2011; Tomonaga et al., 2000) |
| $n_2$ | Cohesin threshold count for the reference chromosome (chromosome 2) | 1 | [0 - 50] | Low number regime reflects that cohesion may persist with few cohesin complexes, consistent with evidence that (i) only a small pool of cohesive cohesin needs to be removed at anaphase (Tomonaga et al., 2000), (ii) chromosome separation is normal with substantially reduced cohesin levels (Heidinger-Pauli et al., 2010), and (iii) that chromosomes fail to separate with substantial cohesin but begin to separate with lower levels of cohesin (Carvalhal et al., 2018). Upper bound allows thresholds that reflect (i) estimates of spindle pulling forces (Grishchuk et al., 2005; Chacón et al., 2014; Gudimchuk and Alexandrova, 2023; Akiyoshi et al., 2010), (ii) the number of microtubules attached to fission yeast kinetochores (Ding et al., 1993), (iii) the force required to mechanically break cohesin (Richeldi et al., 2024), (iv) and potential impacts of cohesion fatigue in response to spindle forces (Sapkota et al., 2018; Daum et al., 2011) . |
| $R_{12}$ | Ratio for the initial cohesin counts of chromosome 1 over chromosome 2 ($N_1/N_2$) | 1 | [0.4 - 2] | Allows chromosome I and chromosome III to have 0.4-2x or 0.5-5x the amount of starting cohesin relative to chromosome II, respectively, based on (i) genome-wide cohesin binding patterns in fission yeast (Schmidt et al., 2009; Mizuguchi et al., 2014), (ii) the fact that centromere organization may influence cohesin levels (Paldi et al., 2020; Yeh et al., 2008), and (iii) assumptions that the size of centromeres may influence cohesin load (Nonaka et al., 2002; Bernard et al., 2001) |
| $R_{32}$ | Ratio for the initial cohesin counts of chromosome 3 over chromosome 2 ($N_3/N_2$) | 1 | [0.5 - 5] | |
| $r_{12}$ | Ratio for the cohesin threshold of chromosome 1 over chromosome 2 ($n_1/n_2$) | 1 | [0.25 - 4] | Range is set to allow up to ~4x difference in effective threshold between chromosomes, based on the observed 2-4 microtubules per kinetochore in fission yeast (Ding et al., 1993; Joglekar et al., 2008) . |
| $r_{32}$ | Ratio for the cohesin threshold of chromosome 3 over chromosome 2 ($n_3/n_2$) | 1 | [0.25 - 4] | |
| $k_{max}$ | Maximum cohesin degradation rate | $sec^{-1}$ | [0.001 - 0.1] | Range spans a large scale of possible maximum rates of cohesin cleavage. |
| $\tau$ | Time to reach $k_{max}$ | sec | [2 - 240]<br><br>[0.5 - 5] for separase autoactivation | Range spans a minutes- to seconds-scale rate of separase activation, to allow gradual activity ramps or rapid separase activation.<br>To mimic separase autoactivation, the range is constrained to seconds-scale. |
| $b$ | Number of cohesin molecules removed per event | 1 | [1 - 50] | Processive separase action model-specific parameter.<br>Range allows small or large "bursts" of separase activity. |
| $n_{inner}$ | Number of cohesin molecules in the innermost core, the | 1 | [1 - 100] | Steric hindrance model-specific parameter. |

| | | | | |
|---|---|---|---|---|
| | region without steric hindrance | | | Range allows a subset of cohesin in the innermost core, where it is not shielded by steric hindrance, but remains well below the upper bound of $N_2$. |
| α | Multiplier modifying the cohesin threshold ($n_2$) | 1 | [0.1 - 0.7] | Constrains MBC treatment conditions to lower $n_2$ range to reflect reduced microtubule forces. |
| $\beta_k$ | Multiplier modifying $k_{max}$ | 1 | [0.1 - 1] | Allows lower $k_{max}$ in separase mutant conditions to reflect reduced separase activity. |
| $\beta_\tau$ | Multiplier modifying the activation timescale (τ) to mimic APC/C mutants | 1 | [1 - 10]<br><br>[1 - 3] for separase autoactivation | Allows higher $\tau$ in APC/C mutant and velcade treatment conditions to reflect slower separase activation. |
| $\beta_{\tau 2}$ | Multiplier modifying the activation timescale (τ) to mimic velcade treatment | 1 | [1 - 20]<br><br>[1 - 3] for separase autoactivation | |

#### Figure S1

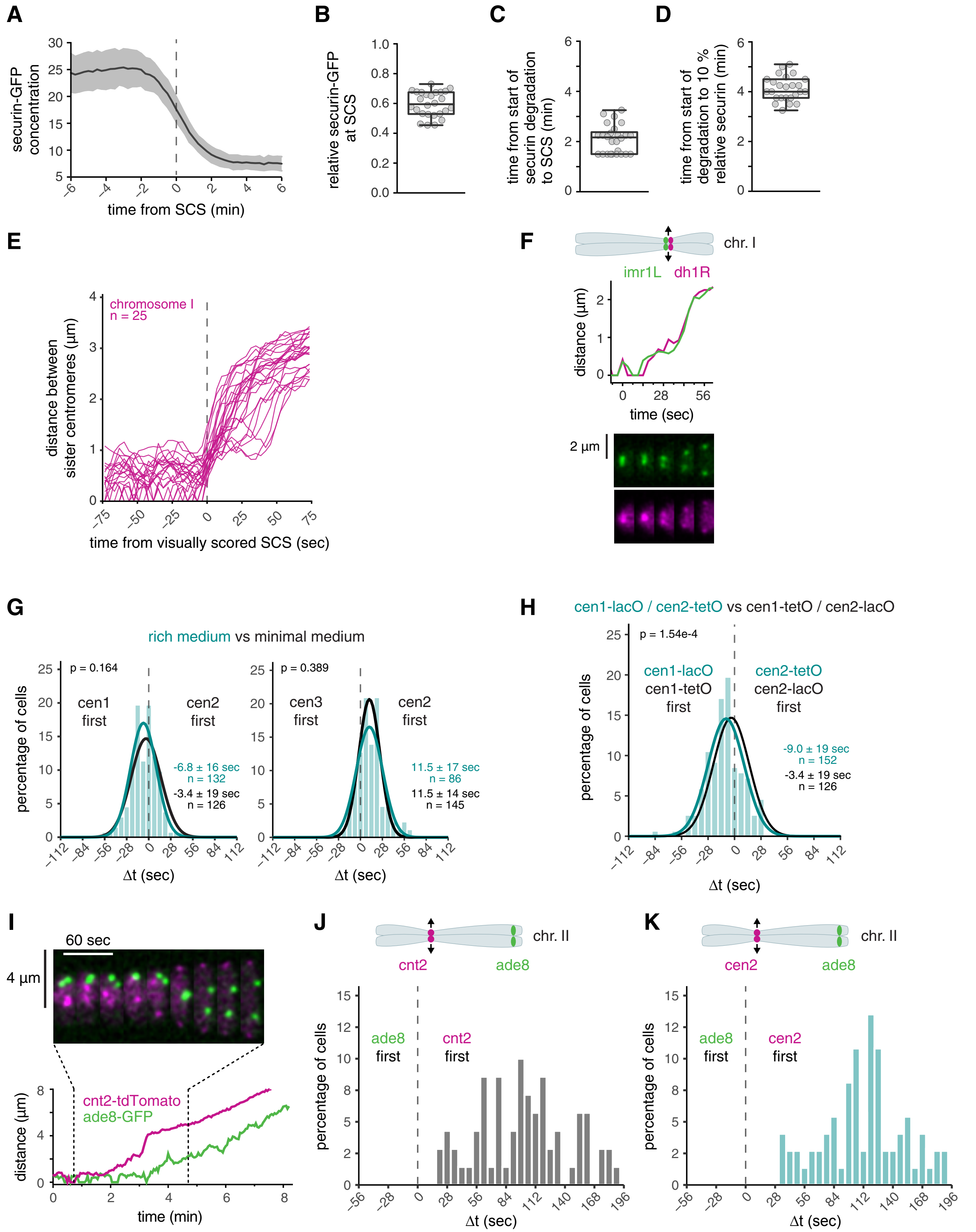

A
securin-GFP concentration
time from SCS (min)
B
relative securin-GFP at SCS
C
time from start of securin degradation to SCS (min)
D
time from start of degradation to 10 % relative securin (min)
E
chromosome I
n = 25
distance between sister centromeres (µm)
time from visually scored SCS (sec)
F
chr. I
imr1L
dh1R
distance (µm)
time (sec)
2 µm
G
rich medium vs minimal medium
p = 0.164
cen1 first
cen2 first
-6.8 ± 16 sec
n = 132
-3.4 ± 19 sec
n = 126
p = 0.389
cen3 first
cen2 first
11.5 ± 17 sec
n = 86
11.5 ± 14 sec
n = 145
percentage of cells
Δt (sec)
H
cen1-lacO / cen2-tetO vs cen1-tetO / cen2-lacO
p = 1.54e-4
cen1-lacO
cen1-tetO
first
cen2-tetO
cen2-lacO
first
-9.0 ± 19 sec
n = 152
-3.4 ± 19 sec
n = 126
percentage of cells
Δt (sec)
I
60 sec
4 µm
cnt2-tdTomato
ade8-GFP
distance (µm)
time (min)
J
chr. II
cnt2
ade8
ade8 first
cnt2 first
percentage of cells
Δt (sec)
K
chr. II
cen2
ade8
ade8 first
cen2 first
percentage of cells
Δt (sec)

**Figure S1. Dynamics of securin degradation and separation of centromeres and chromosome arms.**
**(A)** The nuclear securin-GFP abundance of wild-type cells was followed by live cell imaging (gray, n=26). The cen1-tdTomato marker was used to determine sister chromatid separation (SCS) and individual time courses were aligned to this point (vertical dashed line). Shown is the mean (line) +/- standard deviation (shaded area). **(B)** Quantification of the amount of securin-GFP present at the time of sister chromatid separation (SCS) relative to the amount present at start of degradation. Data is from the experiment shown in (A). Single-cell measurements as circles; the boxplot shows median, interquartile range (box), and range (whiskers). **(C,D)** Quantification of the time from start of securin-GFP degradation until SCS (C) or until 90 % of securin-GFP had been degraded (D). Same representation as described in (B). **(E)** Distances between sister centromeres of chromosome I in a subset of wild-type cells (same experiment as in Fig. 1F). Curves are aligned to sister chromatid separation (SCS) determined by visual inspection of the time-lapse recording. **(F)** Graph of sister chromatid distance and corresponding kymograph for a strain with dh1R-tdTomato and imr1L-GFP markers on the same chromosome. **(G)** Cyan: Frequency distributions and Gaussian fit (continuous lines) of the time difference ($\Delta$t) between the separation of outer centromere markers on chromosome I and II or chromosome II and III for cells grown in rich medium. The fitted Gaussian distributions of $\Delta$t for cells grown in minimal medium are shown in black for comparison. Mean ± standard deviation of the fit; n = number of cells; p-values from a two-sample Kolmogorov-Smirnov test. **(H)** Cyan: Frequency distribution and Gaussian fit (continuous line) of the time difference ($\Delta$t) between the separation of outer centromere markers on chromosome I and II or chromosome II and III using cen1-lacO and cen2-tetO (flipped lacO/tetO relative to the standard tagging). The fitted Gaussian distribution of $\Delta$t for cells with the standard tagging scheme (cen1-tetO, cen2-lacO; same as in Fig. 1F) is shown in black for comparison. Mean ± standard deviation of the fit; n = number of cells; p-values from a two-sample Kolmogorov-Smirnov test. **(I)** Graph of sister chromatid distance and corresponding kymograph for a cell from (J) with chromosome II arm marker (ade8-GFP) and cnt2-tdTomato. **(J,K)** Frequency distributions of the time difference between the separation of a chromosome II arm marker (ade8-GFP) and either cnt2-tdTomato (J) or cen2-tdTomato (K) .

### Figure S2

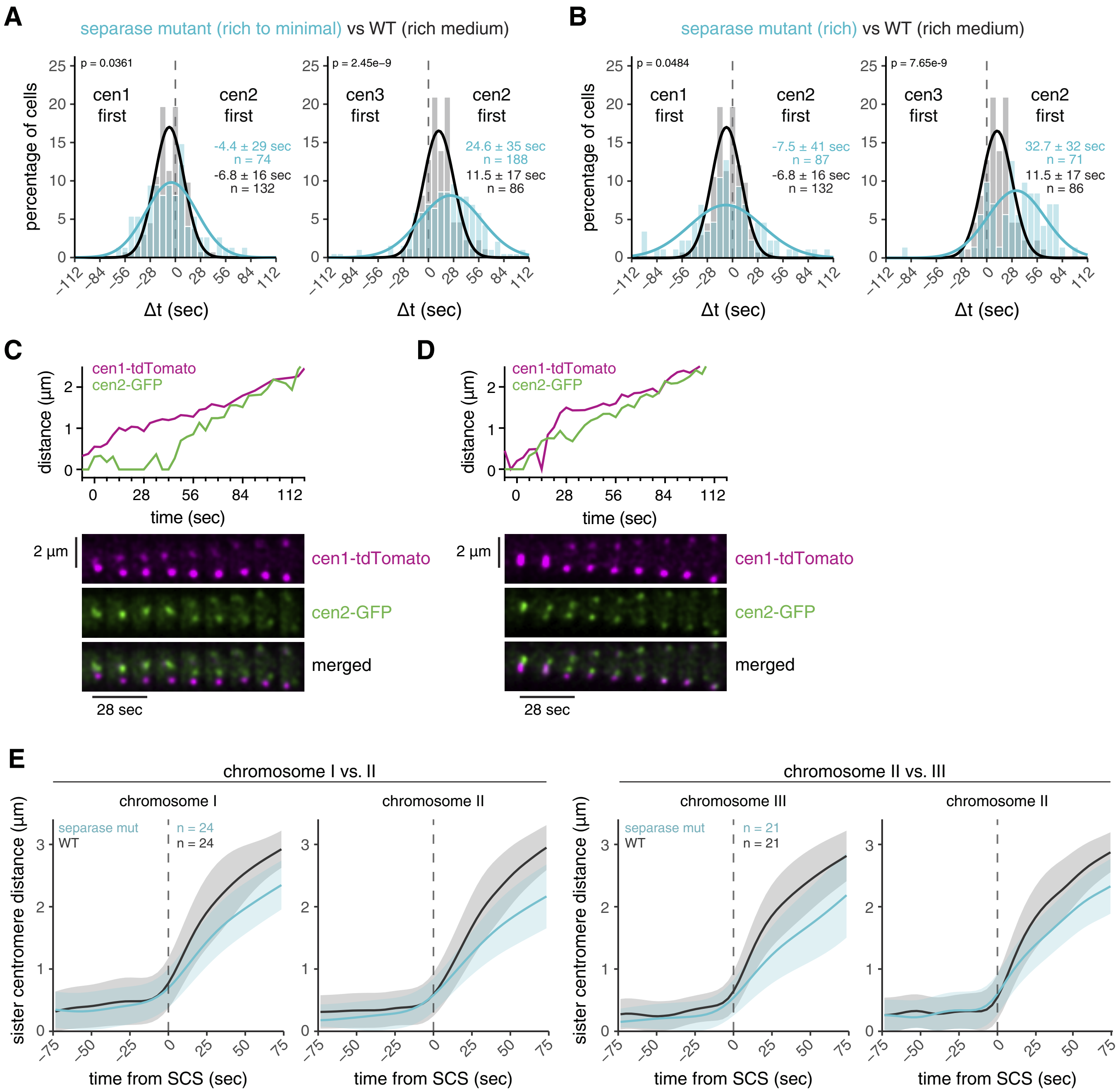

**Figure S2. Lower separase activity reduces centromere separation synchrony and speed.**
**(A,B)** Cyan: frequency distributions and Gaussian fit (continuous lines) of the time difference between the separation of centromere 1 and 2 or centromere 2 and 3 for cells carrying the temperature-sensitive separase allele *cut1-206*, grown in rich medium and then imaged in minimal medium (A) or rich medium (B). The fitted Gaussian distributions of wild-type cells are shown for comparison in black. Mean ± standard deviation of the fit; n = number of cells; p-values from a two-sample Kolmogorov-Smirnov test. Combined separase mutant datasets from (A) and (B) are shown in Fig. 2A, as distributions were not significantly different (Supplementary Table S1). **(C,D)** Sister centromere distance and corresponding kymographs for *cut1-206* cells with cen1-tdTomato and cen2-GFP markers. Examples for asynchronous (C) and more synchronous separation (D). **(E)** Distance between each sister centromere pair in wild-type strains and separase-mutant strains (same cells as shown in Fig. 2B). Distances are aligned to sister chromatid separation (SCS) of the respective chromosome at t=0. Mean is shown as a solid line, standard deviation as shaded area.

#### Figure S3

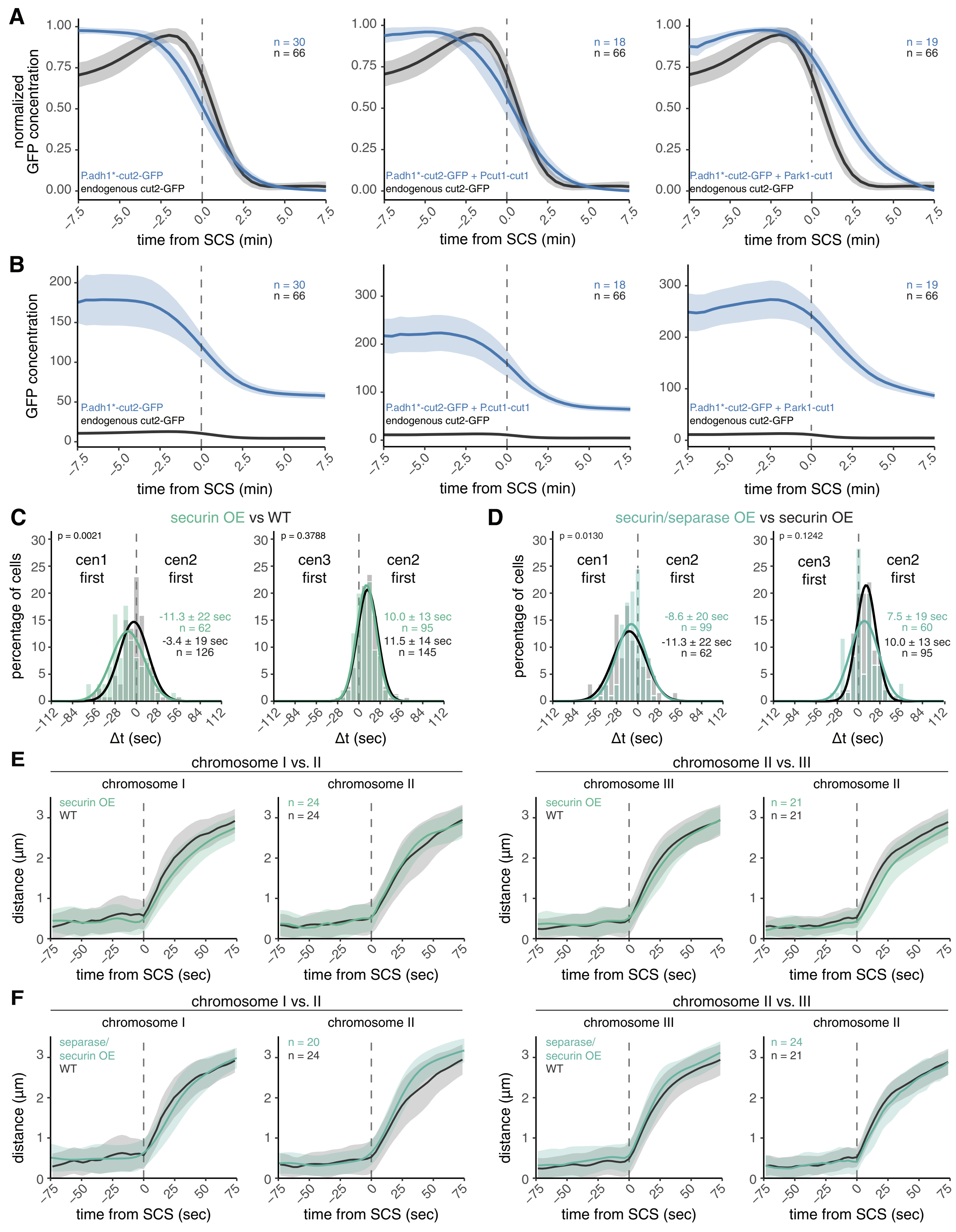

A
normalized GFP concentration
P.adh1*-cut2-GFP
endogenous cut2-GFP
n = 30
n = 66
P.adh1*-cut2-GFP + Pcut1-cut1
n = 18
P.adh1*-cut2-GFP + Park1-cut1
n = 19
time from SCS (min)
B
GFP concentration
P.adh1*-cut2-GFP + P.cut1-cut1
P.adh1*-cut2-GFP + P.ark1-cut1
C
securin OE vs WT
p = 0.0021
cen1 first
cen2 first
-11.3 ± 22 sec
n = 62
-3.4 ± 19 sec
n = 126
p = 0.3788
cen3 first
10.0 ± 13 sec
n = 95
11.5 ± 14 sec
n = 145
percentage of cells
Δt (sec)
D
securin/separase OE vs securin OE
p = 0.0130
-8.6 ± 20 sec
n = 99
-11.3 ± 22 sec
n = 62
p = 0.1242
7.5 ± 19 sec
n = 60
10.0 ± 13 sec
n = 95
E
chromosome I vs. II
chromosome I
chromosome II
securin OE
WT
n = 24
n = 24
chromosome II vs. III
chromosome III
n = 21
n = 21
distance (µm)
time from SCS (sec)
F
separase/ securin OE
n = 20
n = 24
n = 24
n = 21

**Figure S3. Co-overexpression of securin and separase does not strongly change chromosome separation synchrony.** **(A,B)** Normalized (A) and non-normalized (B) concentration measurements of GFP in strains with endogenous cut2-GFP (black) or overexpressing cut2-GFP from the endogenous locus using a mutated adh1 promoter (P.adh1*, blue). Cut2-GFP is either overexpressed alone (left panel), or co-overexpressed with separase, Cut1. For Cut1 overexpression, a second copy was integrated at the leu1 locus, either with the endogenous cut1 promoter (center panel) or the ark1 promoter (right panel). Mean is shown as a solid line, standard deviation as shaded area. **(C,D)** Frequency distributions and Gaussian fit (continuous lines) of the time difference between the separation of centromere 1 and 2 or centromere 2 and 3 for cells with securin overexpression alone (C), or securin co-overexpression with P.cut1-cut1 (D) (same experiment as in Fig. 2C). The fitted Gaussian distributions of wild-type cells (C) or cells with securin overexpression alone (D) are shown for comparison in black. Mean ± standard deviation of the fit; n = number of cells; p-values from a two-sample Kolmogorov-Smirnov test. **(E,F)** Distance between each sister chromatid pair in wild-type cells and cells with securin overexpression (E) or securin and separase co-overexpression (F) (same cells as shown in Fig. 2D). Distances are aligned to the SCS time of the respective chromosome at t=0. Mean is shown as a solid line, standard deviation as shaded area.

#### Figure S4

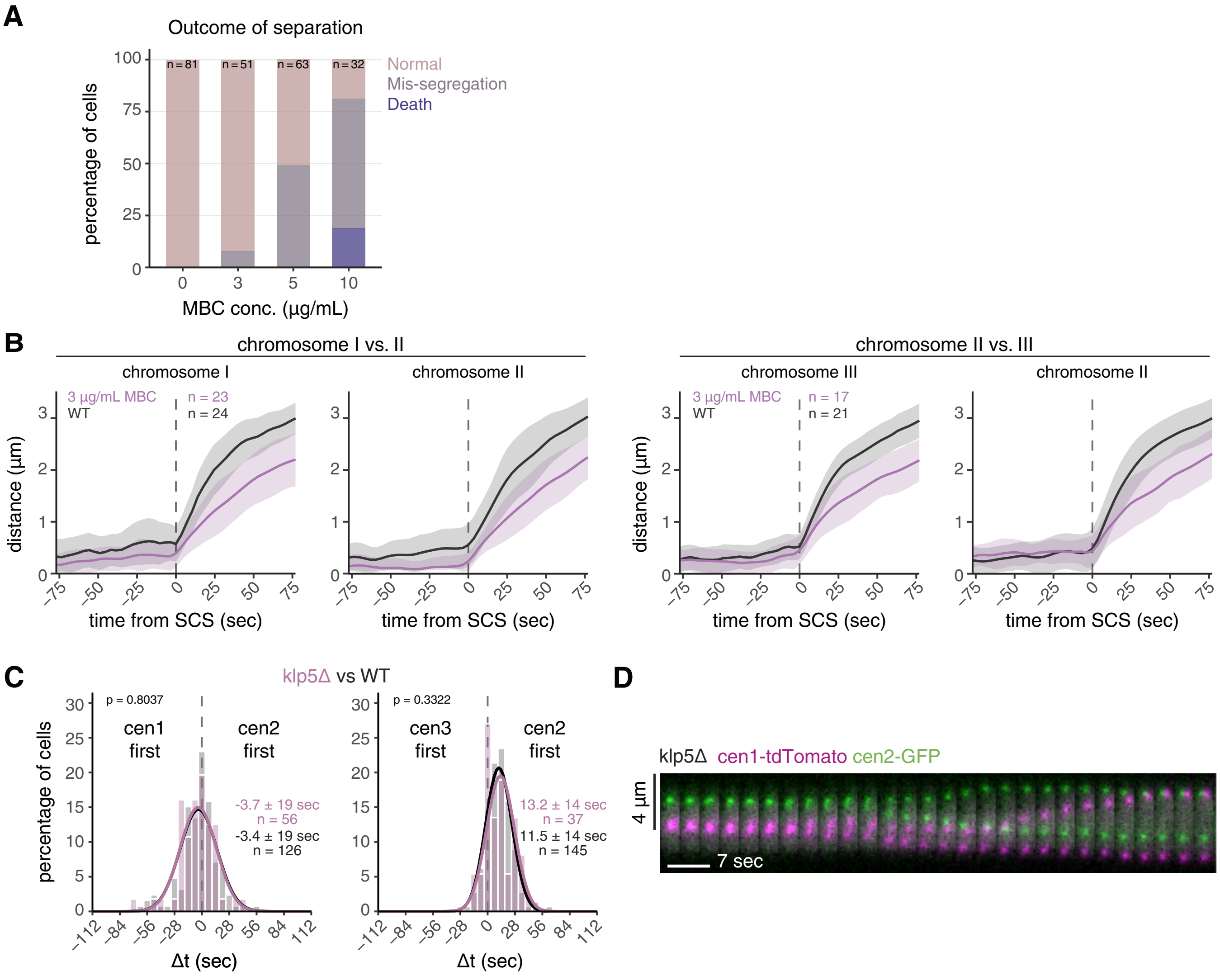

**Figure S4. Influence of MBC and klp5 deletion on sister chromatid separation.**
**(A)** Anaphase outcome of cells which underwent chromosome segregation in medium containing differing concentrations of MBC. Outcomes are shown as a percentage of all cells observed segregating, and were defined as either normal segregation, mis-segregation, or cell death. **(B)** Distance between each sister chromatid pair in wild-type cells and cells with 3 µg/mL MBC (same cells as shown in Fig. 2F). Distances are aligned to the SCS time of the respective chromosome at t=0. Mean is shown as a solid line, standard deviation as shaded area. **(C)** Frequency distributions and Gaussian fit (continuous lines) of the time difference between the separation of centromere 1 and 2 or centromere 2 and 3 for cells with the klp5 gene deleted (klp5Δ). The fitted Gaussian distributions of wild-type cells are shown for comparison in black. Mean ± standard deviation of the fit; n = number of cells; p-values from a two-sample Kolmogorov-Smirnov test. **(D)** Representative kymograph of sister chromatid separation in a klp5Δ cell with cen1-tdTomato and cen2-GFP markers, illustrating the misaligned chromosomes in metaphase.

#### Figure S5

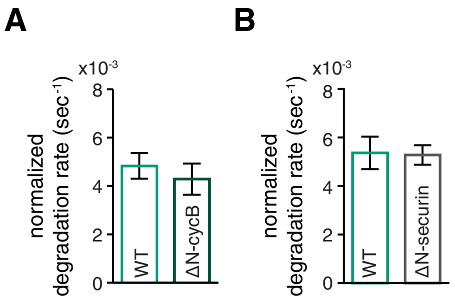

**Figure S5. Securin degradation rate in the presence of non-degradable cyclin B or non-degradable securin.** **(A)** Quantification of the securin degradation rates in the presence (ΔN-cycB, n=22) or absence (WT, n=31) of non-degradable cyclin B (Cdc13) with mean and standard deviation. The corresponding securin degradation curves have been shown previously in Kamenz and Hauf, 2014, Figure 1E. **(B)** Quantification of the securin degradation rates in the presence (ΔN-securin, n=21) or absence (WT, n=27) of non-degradable securin with mean and standard deviation. The corresponding securin degradation curves are shown in Figure 3E.

### Figure S6

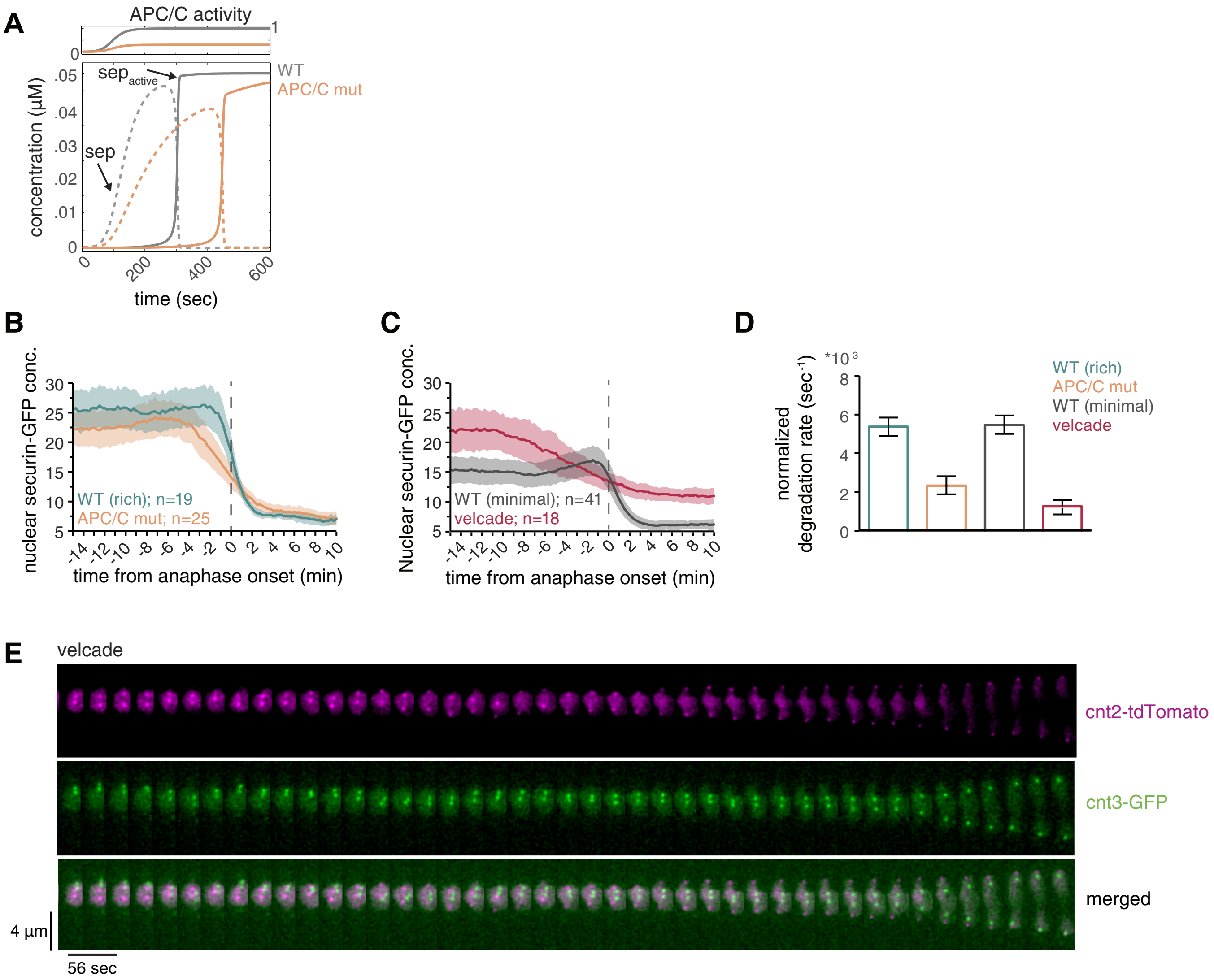

**Figure S6. Modulation of securin degradation kinetics by impairing APC/C or proteasome activity.**
**(A)** Simulation of released separase (dashed lines) and active separase (solid lines) from the model with autocatalytic separase activation (shown in Fig. 4B) assuming high (gray) or low (orange) APC/C activity. See Supplementary Material for details. **(B)** Nuclear securin-GFP intensity was monitored by live-cell imaging of wild-type cells and *cut9-665* temperature-sensitive mutants (APC/C mut). Anaphase onset was determined using the cen1-tdTomato marker, and individual time courses were aligned to this point (vertical dashed line). Shown are the population means (lines) ± standard deviations (shaded areas). **(C)** Nuclear securin-GFP intensity was monitored by live-cell imaging in wild-type cells grown in minimal medium, or cells additionally treated with 100 µM of the proteasome inhibitor velcade (bortezomib). Shown are the population means (lines) ± standard deviations (shaded areas). **(D)** Quantification of the degradation rates for the experiments shown in (B) and (C) after normalization of the data to the minimum and maximum values. Error bars represent the standard deviation. **(E)** Kymograph for a velcade-treated cell with cnt2-tdTomato and cnt3-GFP markers.

### Figure S7

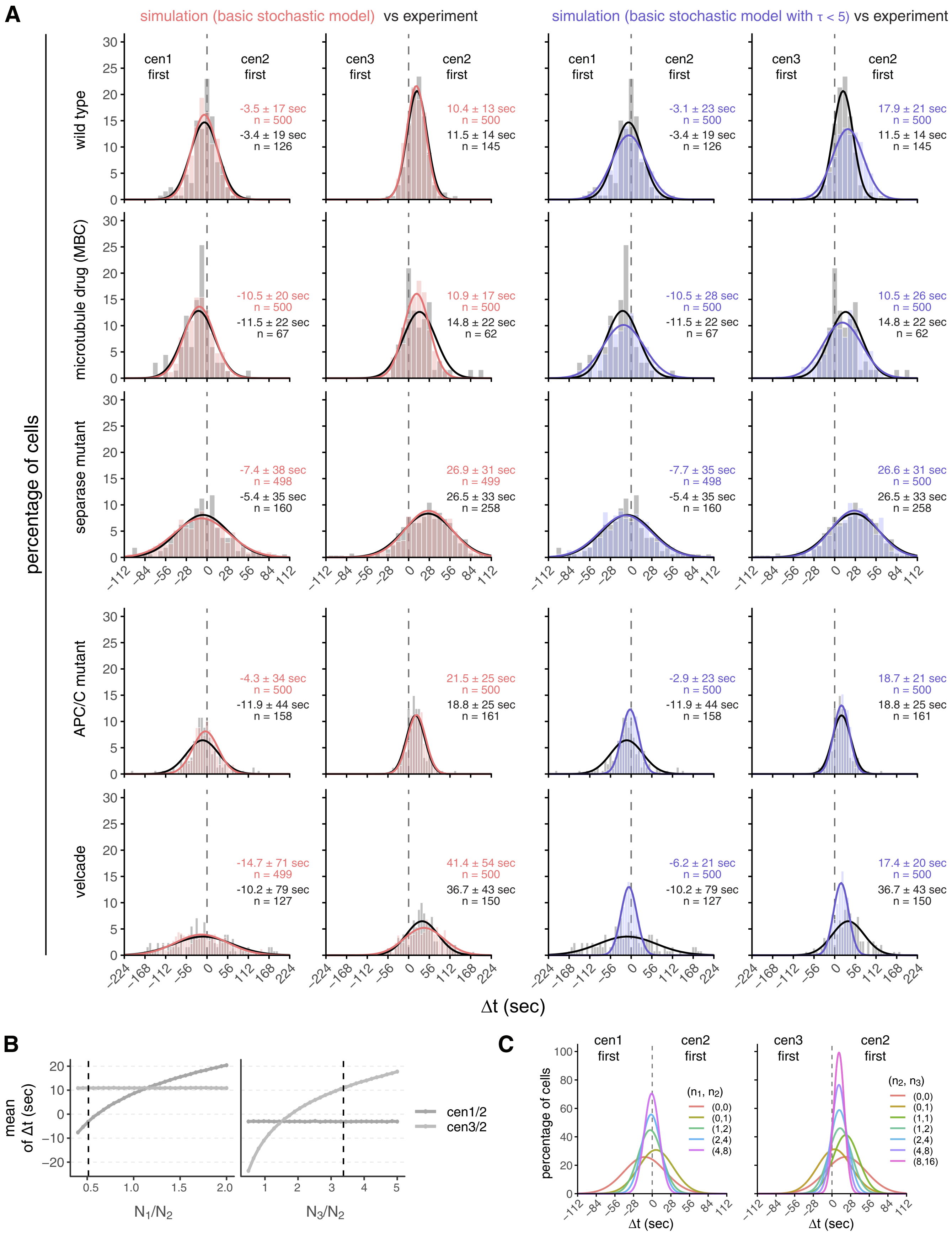
A
simulation (basic stochastic model) vs experiment
simulation (basic stochastic model with τ < 5) vs experiment
percentage of cells
wild type
microtubule drug (MBC)
separase mutant
APC/C mutant
velcade
cen1 first
cen2 first
cen3 first
cen2 first
-3.5 ± 17 sec
n = 500
-3.4 ± 19 sec
n = 126
10.4 ± 13 sec
n = 500
11.5 ± 14 sec
n = 145
-3.1 ± 23 sec
n = 500
-3.4 ± 19 sec
n = 126
17.9 ± 21 sec
n = 500
11.5 ± 14 sec
n = 145
-10.5 ± 20 sec
n = 500
-11.5 ± 22 sec
n = 67
10.9 ± 17 sec
n = 500
14.8 ± 22 sec
n = 62
-10.5 ± 28 sec
n = 500
-11.5 ± 22 sec
n = 67
10.5 ± 26 sec
n = 500
14.8 ± 22 sec
n = 62
-7.4 ± 38 sec
n = 498
-5.4 ± 35 sec
n = 160
26.9 ± 31 sec
n = 499
26.5 ± 33 sec
n = 258
-7.7 ± 35 sec
n = 498
-5.4 ± 35 sec
n = 160
26.6 ± 31 sec
n = 500
26.5 ± 33 sec
n = 258
-4.3 ± 34 sec
n = 500
-11.9 ± 44 sec
n = 158
21.5 ± 25 sec
n = 500
18.8 ± 25 sec
n = 161
-2.9 ± 23 sec
n = 500
-11.9 ± 44 sec
n = 158
18.7 ± 21 sec
n = 500
18.8 ± 25 sec
n = 161
-14.7 ± 71 sec
n = 499
-10.2 ± 79 sec
n = 127
41.4 ± 54 sec
n = 500
36.7 ± 43 sec
n = 150
-6.2 ± 21 sec
n = 500
-10.2 ± 79 sec
n = 127
17.4 ± 20 sec
n = 500
36.7 ± 43 sec
n = 150
Δt (sec)
B
mean of Δt (sec)
cen1/2
cen3/2
N1/N2
N3/N2
C
percentage of cells
cen1 first
cen2 first
cen3 first
cen2 first
(n1, n2)
(0,0)
(0,1)
(1,2)
(2,4)
(4,8)
(n2, n3)
(0,0)
(0,1)
(1,1)
(1,2)
(2,4)
(4,8)
(8,16)
Δt (sec)

**Figure S7. Simulation of sister chromatid separation timing using stochastic models**
**(A)** Experimental data compared to stochastic model simulations of sister chromatid separation times using the basic stochastic model from Fig. 5, or the basic stochastic model with a restriction on the value of $\tau$ ($\tau<5$). Frequency distributions and Gaussian fit (continuous lines) of the time difference ($\Delta t$) between the separation of centromeres 1 and 2 or 2 and 3 are shown for wild-type cells, cells treated with 3 µg/mL MBC, cells carrying the temperature-sensitive separase allele *cut1-206* (separase mutant), cells carrying a temperature-sensitive allele of the APC/C subunit Cut9 (*cut9-665*), and cells exposed to 100 µM velcade. Mean ± standard deviation of the fit; n = number of cells in the experimental data or simulation. **(B)** One-at-a-time (OAT) sensitivity analysis of the basic stochastic model showing the effect of relative cohesin load on mean separation timing. The ratios of initial cohesin numbers between chromosome pairs ($N_1/N_2$ and $N_3/N_2$) were varied individually while all other parameters were held fixed at their optimized values. The resulting mean of the time difference ($\Delta t$) between the separation of centromeres 1 and 2 or 2 and 3 is shown. Dashed vertical lines indicate the optimal parameter values. **(C)** Effect of low cohesin thresholds on separation time variability in the basic model. Gaussian fits of the time difference between the separation of centromeres 1 and 2 or 2 and 3 are shown for simulations in which the separation threshold n was set to specific values for each chromosome (denoted by ($n_1$, $n_2$) or ($n_2$, $n_3$)), while all other parameters were held constant at their optimal values.
